# Global identification of AGO3–RNA interactions in *Chlamydomonas* reveals the small RNA–mediated regulation of nuclear and chloroplast gene expression

**DOI:** 10.1101/2024.10.23.619964

**Authors:** Suzuna Murakami, Hiroki Takahashi, Kaede Shimizu, Tomohito Yamasaki

## Abstract

MicroRNAs (miRNAs) form complexes with Argonaute (AGO) proteins and bind to mRNAs with complementary sequences to repress their expression. Organisms typically possess several hundred miRNAs regulating diverse biological processes. Although the roles of miRNAs have been well elucidated in multicellular organisms, they remain largely unexplored in unicellular organisms. In the green alga *Chlamydomonas reinhardtii*, the first unicellular organism in which miRNAs were discovered, identifying miRNA target genes remains challenging. Here, we report the high-throughput sequencing of RNA isolated by crosslinking immunoprecipitation (HITS-CLIP) to generate global AGO3–RNA interaction maps, identifying 131 mRNAs derived from nuclear genes. We also detected mRNAs from two chloroplast genes, suggesting that AGO3 may also influence chloroplast gene expression. Our findings indicate that *Chlamydomonas* small RNAs repress nuclear and chloroplast gene expression. Furthermore, HITS-CLIP analyses are now feasible for any RNA-binding protein in *Chlamydomonas*, offering new opportunities to uncover the functions of RNA-binding proteins of interest.

## Introduction

Small RNAs (sRNAs), such as 20–25-nucleotide (nt) microRNAs (miRNAs), associate with Argonaute (AGO) proteins to form the RNA-induced silencing complex (RISC), which binds to complementary messenger RNAs (mRNAs) to repress their expression. In animals, complementarity between the target transcript and the seed region (2–8 nt at the 5’ end of the miRNA) is key for binding^1–3^. The repression of expression can occur via translation inhibition, mRNA decapping, mRNA de-adenylation, and/or mRNA destabilization^4–7^. There are also a few documented cases of miRNA-directed mRNA cleavage in mammals^8–10^. In plants, miRNAs generally require near-perfect complementarity to their target mRNAs, leading to mRNA cleavage or translation repression and thus to effective expression repression^11,12^.

The unicellular green alga *Chlamydomonas reinhardtii* has a functional sRNA-mediated silencing system involving AGO3, the double-stranded RNA–specific ribonuclease Dicer-like3 (DCL3) and the double-stranded RNA–binding protein Dull slicer-16 (DUS16)^13–15^. Although sRNAs regulate many aspects of biology in multicellular organisms, their roles in *Chlamydomonas* are less understood^16^, partly due to the poor conservation between algal miRNAs and those of plants or animals^17–19^. Furthermore, single *ago3*, *dcl3*, or *dus16* mutants show minimal phenotypic changes, complicating their functional analysis^13–15,20^. Identifying the target genes of *Chlamydomonas* miRNAs is therefore important for understanding their roles.

Earlier attempts to identify *Chlamydomonas* miRNA target genes through computational methods, including predictions of miRNA–mRNA complementarity, transcriptome deep sequencing (RNA-seq), and ribosome footprinting (Ribo-seq), have had limited success^20–22^. Similar challenges have been noted in animals, but their miRNA targets can be reliably identified using crosslinking immunoprecipitation followed by high-throughput sequencing (CLIP-seq), a genome-wide technique for mapping RNAs associated with RNA-binding proteins (RBPs)^23^. Among the CLIP-seq methods, the high-throughput sequencing of RNA isolated by crosslinking immunoprecipitation (HITS-CLIP) stands out^24^. In this technique, RBPs are crosslinked to the RNAs they bind in living cells using ultraviolet (UV) light, the crosslinked RNA is partially digested with RNase, and then the RBP–RNA complexes are immunopurified for library preparation and sequencing. The binding footprints of the RBP are identified as read clusters on the reference genome and gene models^25^. Additionally, reverse transcription errors introduced at crosslinking sites, known as crosslink-induced mutation sites (CIMS), help pinpoint RBP-binding regions when analyzed with bioinformatics tools such as the CTK Toolkit^25,26^. Overlapping CIMS and read clusters reliably indicate RBP-binding sites.

Here, we developed a HITS-CLIP pipeline for AGO3 by adapting a published protocol for cultured mammalian cells^25^ with modifications tailored for *Chlamydomonas*. We identified 133 candidate transcripts derived from nuclear and chloroplast genes that are bound by AGO3–RISC, indicating that sRNAs are likely to play diverse regulatory roles even in the unicellular *Chlamydomonas*.

## Results

### Optimization of the HITS-CLIP technique for *Chlamydomonas* AGO3

We used a *Chlamydomonas* complementation strain expressing *AGO3* with the sequence for a FLAG tag in the *ago3-1* mutant background^13^. We constructed the AGO3 HITS-CLIP libraries according to the pipeline illustrated in Supplementary Fig. 1 (see Methods for details). In this method, we used ∼254 nm UV light to induce irreversible crosslinking between the RNA and proteins, optimizing the UV energy to the RNA-binding protein, in this case AGO3. We then fragmented the mRNA crosslinked to AGO3 to ∼50 nt in length to identify AGO3-binding regions, as it has been reported that the footprint of mouse AGO2 on mRNA is ∼46 nt, with 50 nt being the optimal size for bioinformatic analysis^25,27^. To assess the extent of AGO3–RNA crosslinking obtained with various UV energy levels, we analyzed samples irradiated with varying UV doses using a northwestern analysis. We thus determined that 1,000 mJ/cm² UV is the optimal dose for AGO3 (Supplementary Fig. 2a). RNase A optimization is necessary to fragment the RNA to ∼50 nt while preventing over-digestion, even when the RNA is bound to AGO3. We then incubated the crosslinked material was treated with different RNase A concentrations, looking for an appropriate smear above the AGO3 band, reflective of partial degradation of the RNA molecules to ∼50 nt. We chose a 1/5,000 dilution (0.5 ng/ml) for RNase A (Supplementary Fig. 2b). Using the optimized conditions above (see Methods), we obtained read lengths of the desired range for the sRNA and target RNA (Supplementary Fig. 2c, d).

### AGO3 HITS-CLIP successfully identifies known AGO3-bound sRNA sequences

We collected cells cultured under mixotrophic (with acetate as a reduced carbon source) or phototrophic conditions to generate HITS-CLIP libraries for AGO3-bound sRNA and AGO3-bound fragmented mRNA, termed sRNA and target libraries hereafter (Supplementary Fig. 1). We processed all raw reads into deduplicated reads for downstream informatics analysis (Supplementary Fig. 3, Supplementary Table 1). In line with previously reported AGO3-associated sRNAs^21^, most sRNA reads were 21 nt in length, with a T (corresponding to U in RNA) at their 5’ ends (Fig. 1).

**Fig. 1.**
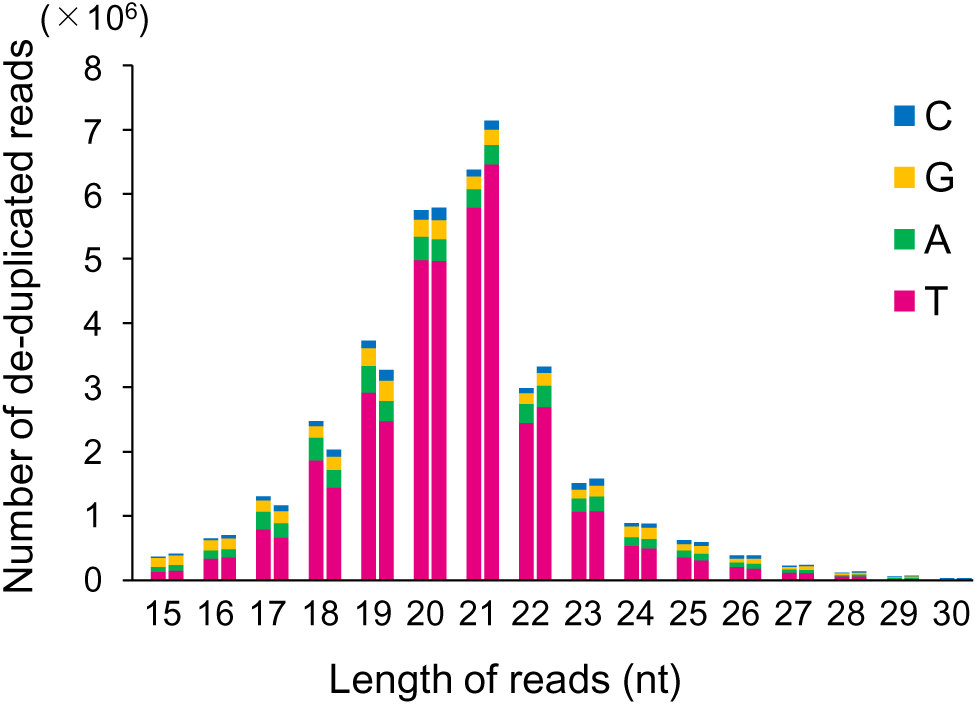
The peak length of the sRNA read distribution is 21 nt, with the 5’ end of most reads starting with T. For each read length between 15 and 30 nt, the two bars indicate the data for sRNAs under mixotrophic conditions (left) or phototrophic conditions (right). nt, nucleotides.

To verify that these sRNA sequences were dependent on crosslinking, we looked for CIMS and counted the number of deletion, substitution, and insertion mutations (globally designated as *m*) at each position across all unique tags (with *k* representing the total number of tags covering the mutation site). From these values, we calculated the mutation frequency *m*/*k* (Supplementary Fig. 3). A CIMS analysis of 20–22-nt unique tags revealed that deletion-type CIMS cluster primarily in the regions corresponding to sRNAs in the genome (Supplementary Table 2). By focusing on abundant CIMS with *k* ≥ 100 and excluding those likely to reflect sequence polymorphisms between the wild-type strain used here and the *Chlamydomonas* reference genome (based on *m*/*k* ≥ 0.90), we successfully annotated 90% of the remaining deletion-type CIMS to specific sRNA loci^28^ (Supplementary Table 2). Substitution-type CIMS were also well annotated but were less frequent than the deletion-type CIMS. By contrast, the rates of insertion-type CIMS were very low at sRNA loci, indicating that insertions at crosslinking sites are rare. These findings suggest that most CIMS at AGO3–sRNA crosslinking sites are deletions or substitutions, consistent with previous AGO2 HITS-CLIP results obtained for mouse (*Mus musculus*)^29^. Deletions and insertions were primarily located near the central region of the sRNA sequence, while substitution-type CIMS followed a largely constant distribution over the length of the sRNA (Supplementary Fig. 4), mirroring the results of the previous mouse AGO2 analysis^29^.

We detected clusters of unique tags and CIMS mapping to *MIRNA* loci, the mature miRNAs of which are known to be associated with AGO3. The previous HITS-CLIP analysis of mouse AGO2 data for deletion-type CIMS with *m*/*k* ≥ 0.08 within these clusters were considered to be AGO2-binding regions^29^. When we analyzed 20–22-nt sRNA unique tags against the entire *Chlamydomonas* genome reference, we observed that these unique tags form clusters at most *MIRNA* loci, with deletion-type and/or substitution-type CIMS (*m*/*k* ≥ 0.08) identified within these clusters. The *MIR912* locus is shown in Fig. 2, while *MIR1153* and *MIR1157* are shown in Supplementary Fig. 5. These results clearly indicate that AGO3 HITS-CLIP successfully identified AGO3-associated sRNAs in *Chlamydomonas*.

**Fig. 2.**
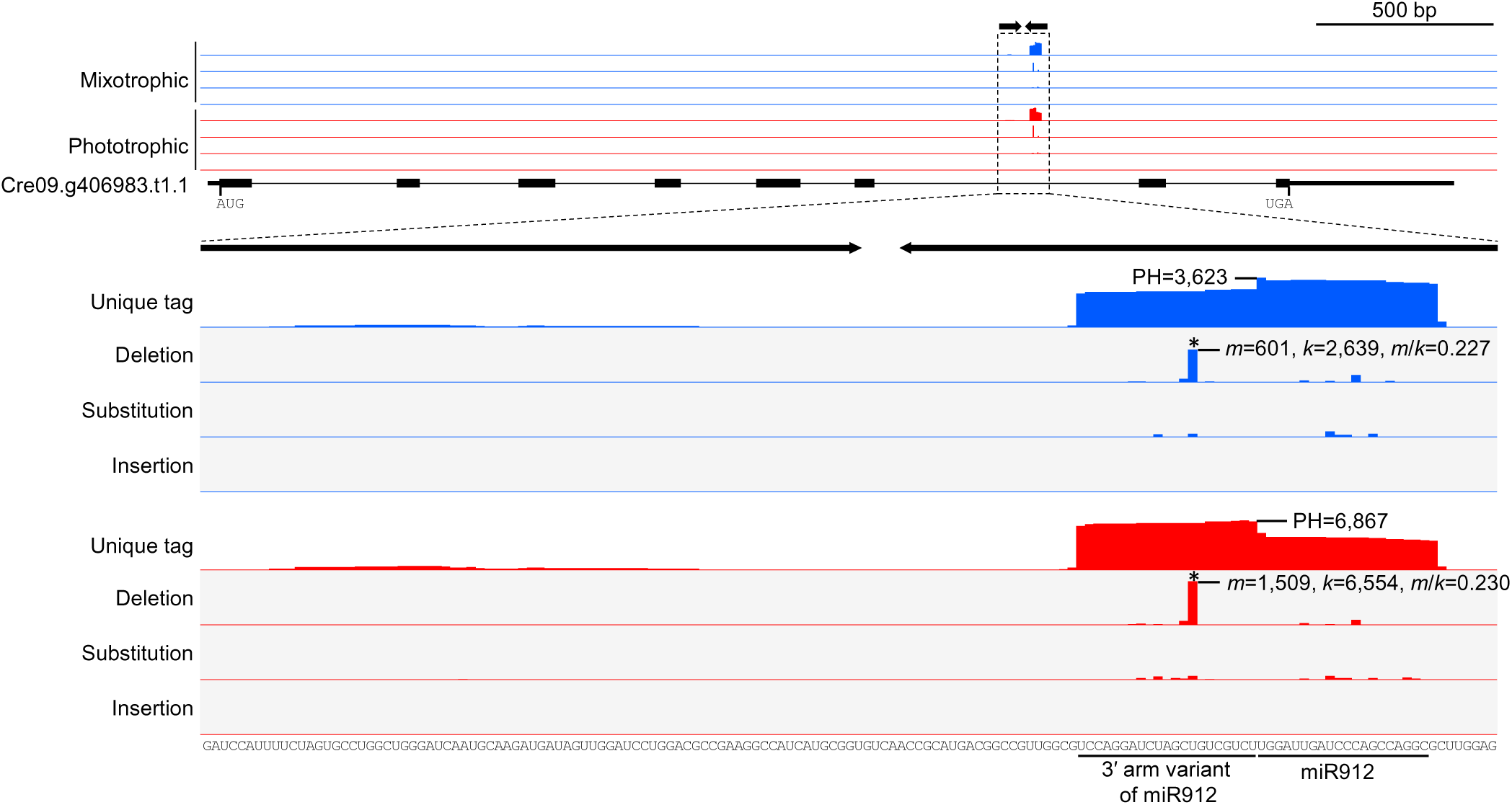
Cluster of unique sRNA tags and location of crosslink-induced mutation sites were found at the *MIR912* locus. For each condition, the panels show (top to bottom) the coverage of unique tags and the positions of deletions, substitutions, and insertions detected through crosslink-induced mutation site (CIMS) analysis. The *x*-axis shows either the coverage of unique tags or the proportion of CIMS (*m*/*k*). The top panel provides an overview of the pre-mRNA form of Cre09.g406983.t1.1, where the primary transcript of *MIR912* is located in the 6th intron. The inverted repeat region corresponding to the primary *MIR912* transcript (*pri-MIR912*) is indicated as opposing arrows and highlighted with a dashed box. An enlarged view of this region is presented in the lower panel. The sequence of this region is shown below, with the sequence of the mature miRNA underlined. PH, peak height; *m*, number of mutations; *k*, number of unique tags covering the mutation site. Asterisk (*) marks CIMS with *m*/*k* ≥ 0.08.

### Successful detection of artificial target mRNA cleaved by AGO3–RISC

The peak length of the target reads in the distribution was 47 nt (Supplementary Fig 2), which is nearly identical to the footprint size of mouse AGO2 on mRNA (∼46 nt)^23^. This suggests that the reads in the library were primarily derived from mRNA bound to *Chlamydomonas* AGO3. We analyzed reads 30–100 nt in length from the target library (Supplementary Table 1) to map genes whose transcripts may be targeted by AGO3–RISC. The strain used in generating sequencing libraries harbors a *Gaussia* luciferase reporter construct containing a target site for miR9897-3p in the 3’ untranslated region (3’ UTR) of the reporter, leading to the mRNA cleavage of the artificial *Gluc* mRNA by FLAG-AGO3, and thus a lower Gluc activity^13,30^. We detected a cluster of unique tags mapping to the reporter sequence, which appear to be truncated mRNAs downstream of the miR9897-3p cleavage site (Fig. 3). We did not detect clusters upstream of the cleavage site. Even the upstream fragment is detectable via a northern blot analysis^13^, likely because their 3’ ends were masked by AGO3, hindering the ligation of the 3’ adaptor ligation. We identified several cases of deletion-type and substitution-type CIMS across the entire length of the tag cluster and thus not restricted to the miR9897-3p recognition sequence (Fig. 3). The highest *m*/*k* ratio for a deletion-type CIMS was 0.022 and that for a substitution-type CIMS was 0.036, both below the 0.08 significance threshold for reliable identification previously applied in the mouse AGO2 study. This observation suggests that unstable binding between AGO3 and the *Gaussia* luciferase mRNA truncated at its 5’ end, possibly affecting crosslinking efficiency. These results demonstrate that unique tags from the target library cluster around miRNA target sites, with both deletion and substitution changes detected within the target sequences.

**Fig. 3.**
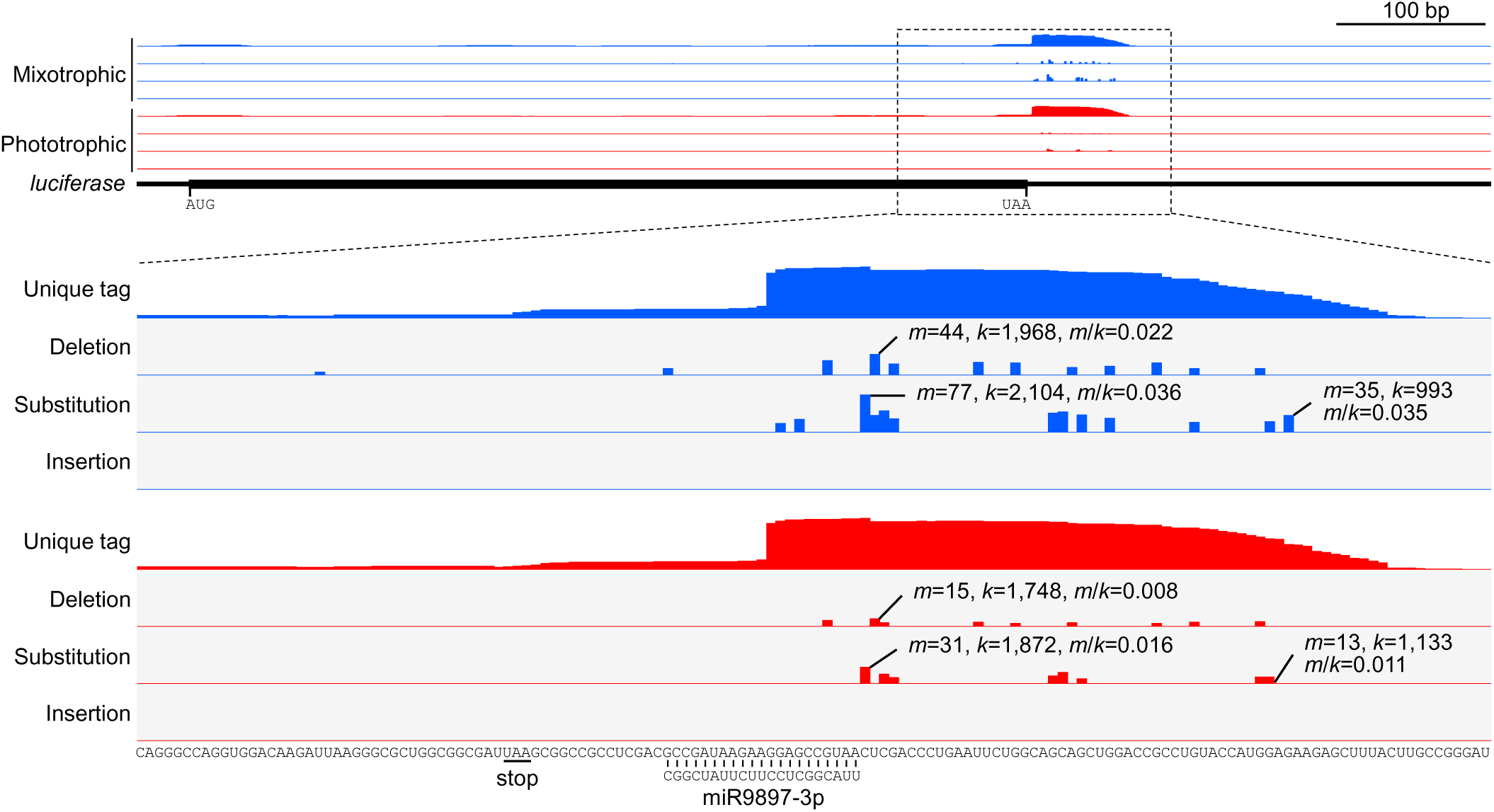
Unique tag cluster and CIMSs were found on miR9897-3p-cleaved luciferase mRNA sequences. The top panel provides an overview of the *Gaussia luciferase* transgene. The sequence of miR9897-3p, complementary to the engineered target, is displayed in a 3’-to-5’ orientation beneath the expanded region sequences.

### Multiple criteria were established to exclude false positives, resulting in the detection of 131 candidate AGO3-binding endogenous transcripts

The presence of deletion-type and substitution-type CIMS, along with clusters of unique tags, indicate mRNAs that bind to AGO3; however, we were concerned that the total number of detected CIMS and clusters, despite a false discovery rate (FDR) < 0.001, may still include many false positives.

The distribution of *m*/*k* values revealed some deletion-type and substitution-type CIMS above 0.08 (Supplementary Table 3). While *m*/*k* values for CIMS mapping to the luciferase mRNA were well below 0.08 (the highest value being 0.036), we set *m*/*k* ≥ 0.08 as a high-confidence criterion, anticipating higher *m*/*k* values for AGO3 targets subjected to translation inhibition. Moreover, single-nucleotide polymorphisms (SNPs) are present between the genomes of the Gluc(1x) strain and the reference *Chlamydomonas* strain; these positions will appear as CIMS with a *m*/*k* value close to 1, prompting us to exclude CIMS with *m*/*k* ≥ 0.95. Additionally, we set *k* ≥ 100 as a minimal coverage criterion. Applying these two stringent criteria, we identified 426 deletion-type CIMS mapping to 283 transcripts and 262 substitution-type CIMS mapping to 137 transcripts under mixotrophic conditions. Under phototrophic conditions, we identified 233 deletion-type CIMS mapping to 134 transcripts and 185 substitution-type CIMS mapping to 82 transcripts (Supplementary Table 4). We noticed no clear bias in the location of these high-confidence CIMS (Supplementary Table 5).

The peaks of these high-confidence unique tag clusters showed high peak height (PH) and low background (PH0) values (Supplementary Fig. 3). Based on the distribution of PH and PH0/PH values, we selected high-confidence peaks using the criteria PH ≥ 100 and PH0/PH ≤ 0.1 (Supplementary Table 6). We thus identified 463 high-confidence peaks mapping to 434 transcripts under mixotrophic conditions and 114 high-confidence peaks mapping to 107 transcripts under phototrophic conditions (Supplementary Table 7). As for CIMS, we observed no clear bias in the position of the selected high-confidence peaks along the mRNAs (Supplementary Table 8).

We defined the region covered by each unique tag cluster as half of the PH (halfPH) with a 30-nt extension on either side of the high-confidence peaks (Supplementary Fig. 3). We then looked for peaks with high-confidence CIMS within this range, using the presence of CIMS as an indication of true crosslinking. Under mixotrophic conditions, we detected high-confidence deletion-type and/or substitution-type CIMS in the halfPH ± 30-nt region of 130 peaks mapping to 129 transcripts, with 12 peaks showing both types of CIMS mapping to 12 transcripts (Supplementary Table 9). Similarly, under phototrophic conditions, we detected high-confidence deletion-type and/or substitution-type CIMS in the halfPH ± 30-nt region of 37 peaks mapping to 36 transcripts, with two peaks showing both types of CIMS mapping to two transcripts (Supplementary Table 10).

Using the halfPH ± 30-nt sequences as high-confidence peaks for queries, we looked for potential sRNA binding sites by conducting an analysis with the online tool psRNAtarget. To this end, we prepared an sRNA input file by selecting those sRNAs detected from the redundant reads (20–22 nt) with a perfect match to the *Chlamydomonas* genome (version 6, https://phytozome-next.jgi.doe.gov/info/CreinhardtiiCC_4532_v6_1), thus excluding those with CIMS. Additionally, we only retained relatively abundant sequences supported by at least 100 reads per million, resulting in 308 sRNA sequences from mixotrophic conditions (Supplementary Table 11) and 376 from phototrophic conditions (Supplementary Table 12). As input files for the target sequences, we used the above high-confidence peak halfPH ± 30-nt regions (Supplementary Tables 9 and 10). We adjusted the parameters of the psRNAtarget analysis to emphasize seed sequence complementarity (FASTA files for the analysis provided in Supplementary Data 1). This analysis identified 600 sRNA–target pairs across 117 transcripts under mixotrophic conditions (Supplementary Table 13) and 217 sRNA–target pairs across 35 transcripts in phototrophic conditions (Supplementary Table 14). In total, we identified 131 transcripts as potential AGO3–RISC targets, with 20 transcripts detected under both conditions (Supplementary Table 15).

### Expression analysis of *CAS*, a newly identified AGO3 target

Cre12.g497300 encodes the chloroplast calcium sensor protein CAS^31^. We detected unique tag clusters mapping to Cre12.g497300.t1.2, with high-confidence deletion-type CIMS with an *m*/*k* value of 0.207 within the cluster coverage under mixotrophic conditions (Fig. 4a, Supplementary Tables 11 and 12). We observed no change in CAS protein abundance between the wild type and the *ago3-1* mutant grown under non-synchronized culture conditions, but CAS abundance was higher in the mutant in the synchronized cultures (Fig. 4b). The CAS protein levels largely returned to wild-type levels in the *ago3-1* C complementation strain (Fig. 4c), and the *CAS* mRNA levels were partially restored to the wild-type levels (Fig. 4d). We observed a similar trend for the *CAS* mRNA and CAS protein levels in the *dus16-1* mutant (Supplementary Fig. 6), with slightly shorter poly(A) tails observed in the mutant relative to the wild type (Supplementary Fig. 7). These results indicate that AGO3–RISC represses *CAS* expression, confirming that AGO3 HITS-CLIP can successfully identify target genes. The elevated levels of *CAS* mRNA in the mutants suggest that AGO3–RISC can decrease target mRNA levels through mechanisms that are independent of slicer activity.

**Fig. 4.**
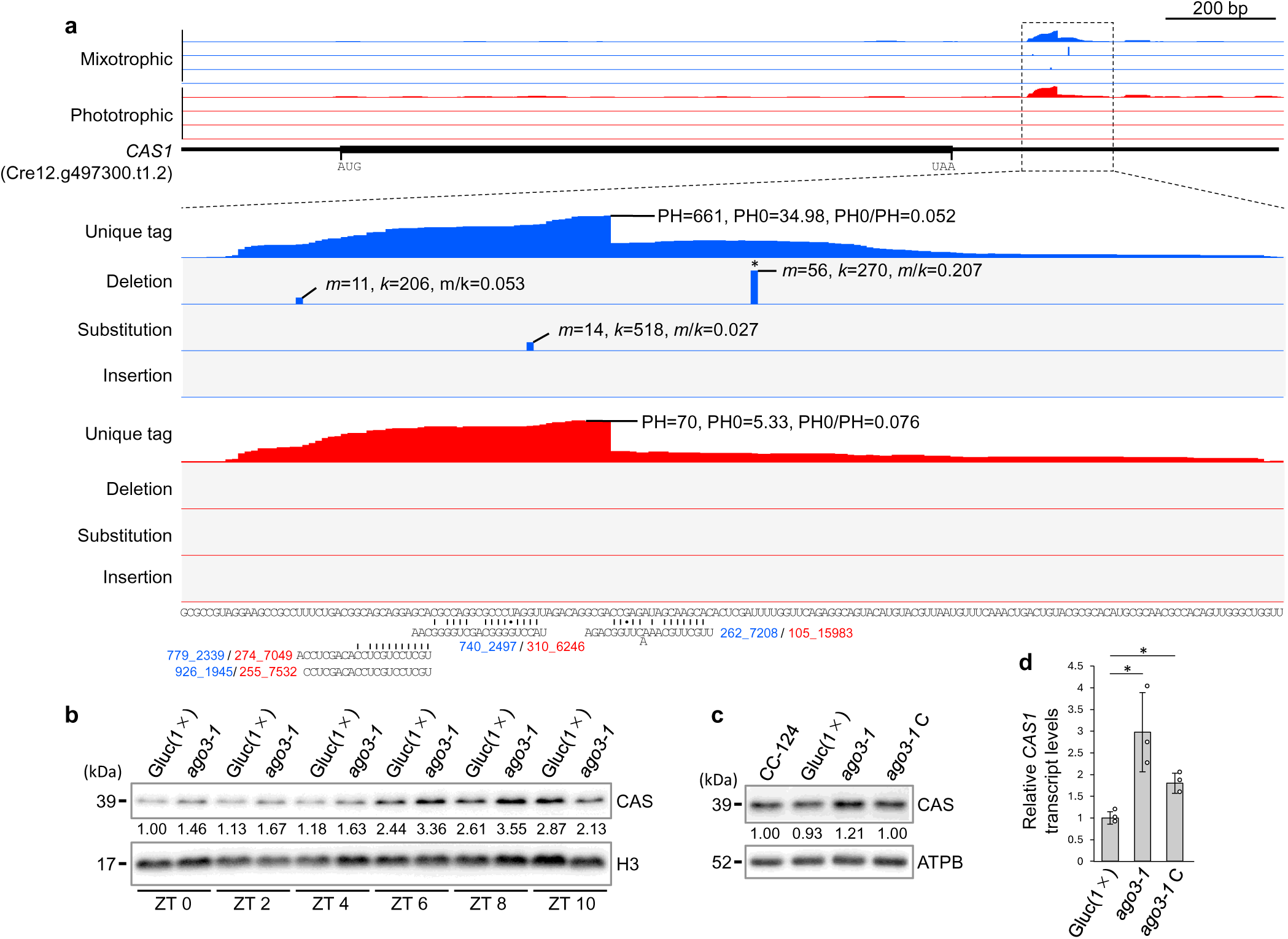
AGO3–RISC binds to the 3’ UTR of *CAS* transcripts, and increased *CAS* protein levels are present in the *ago3* mutant. (a) Location of unique tag clusters and CIMS on the *CAS* transcript (Cre12.g497300.t1.2). The sequences of sRNAs that can recognize the *CAS* transcript are shown in a 3’-to-5’ orientation at the bottom, with corresponding the sRNA IDs (blue for mixotrophic conditions, red for phototrophic conditions) listed in Supplementary Tables 11 and 12. (b) Immunoblot analysis of CAS protein abundance in the Gluc(1×) and *ago3-1* strains. Gluc(1×) is the parental strain of the *ago3-1* mutant. Both strains were grown under a 12-h light/12-h dark photoperiod to synchronize their cell division, and protein samples were collected every 2 h from the start of the light period to 10 h (zeitgeber time 0 (ZT0) to ZT10). Histone H3 was used as a loading control. CAS and H3 protein levels were quantified from the signal intensity, and CAS abundance was normalized to that of H3, with the values from Gluc(1×) set to 1 at ZT0. (c) Immunoblot analysis of CAS protein abundance in the *ago3-1* genetic complementation strain. CC-124 is the parental strain of Gluc(1×), and *ago3-1* C is the partly complemented strain expressing *FLAG-AGO3* in the *ago3-1* background, which was used for the HITS-CLIP analysis. Protein samples were taken at ZT8, with ATPB used as a loading control. CAS and ATPB protein levels were quantified from the signal intensity, and CAS abundance was normalized to that of ATPB, with the values from CC-124 set to 1. (d) RT-qPCR analysis of *CAS* mRNA abundance in the indicated strains. RNA samples were collected from synchronized cells at ZT6. *CAS* transcript levels were normalized to those of *NUOP4*, an *NADH* oxidoreductase gene that exhibits consistent expression throughout the cell cycle. The value corresponding to Gluc(1×) for *CAS* levels was set to 1, and the other values were adjusted accordingly. The values presented are the means ± SD of three independent biological replicates. A two-tailed, unpaired t-test was employed to identify statistically significant differences. Asterisks indicate statistically significant differences (*P* < 0.05).

In addition to the unique tag clusters mapping to the *CAS* locus, we noticed an asymmetric pattern in the clusters, with a cliff of unique tag clusters on the 3’ side for some transcripts (Supplementary Table 15). The presence of these cliffs suggests a hotspot on the 3’ side of the fragmented mRNA that binds to AGO3, although its relevance to AGO3 binding (e.g., footprint) is not known.

Notably, we identified *FtsZ2*, which encodes a chloroplast division ring protein, and *CCS1*, involved in cytochrome *c* biosynthesis, as potential AGO3–RISC target candidates, although we did not test the abundance of their corresponding proteins in the wild type and the *ago3-1* mutant (Supplementary Fig. 8, Supplementary Table 15). If AGO3–RISC represses the expression of these genes, the sRNAs likely regulate various aspects of chloroplast physiology.

### AGO3–RISC may act on chloroplast RNAs

While the sRNA analysis did not identify high-confidence clusters mapping to the genome of either organelle, we did reveal several high-confidence peaks (PH ≥ 100, PH0/PH ≤ 0.1) among the target reads mapping to the chloroplast genome. Deletion-type and/or substitution-type CIMS were present within the halfPH ± 30-nt region of these peaks in the 5’ UTRs of the *petA* and *psaC* loci (Fig. 5, Supplementary Fig. 9). An RT-qPCR and immunoblot analysis showed that *petA* mRNA and cytochrome *f* (cyt*f*) protein levels are higher in the *ago3-1* and *dus16-1* mutants (Fig. 5). Attempts to analyze PsaC protein levels yielded inconsistent results due to its small size (∼9 kDa) and high abundance.

**Fig. 5.**
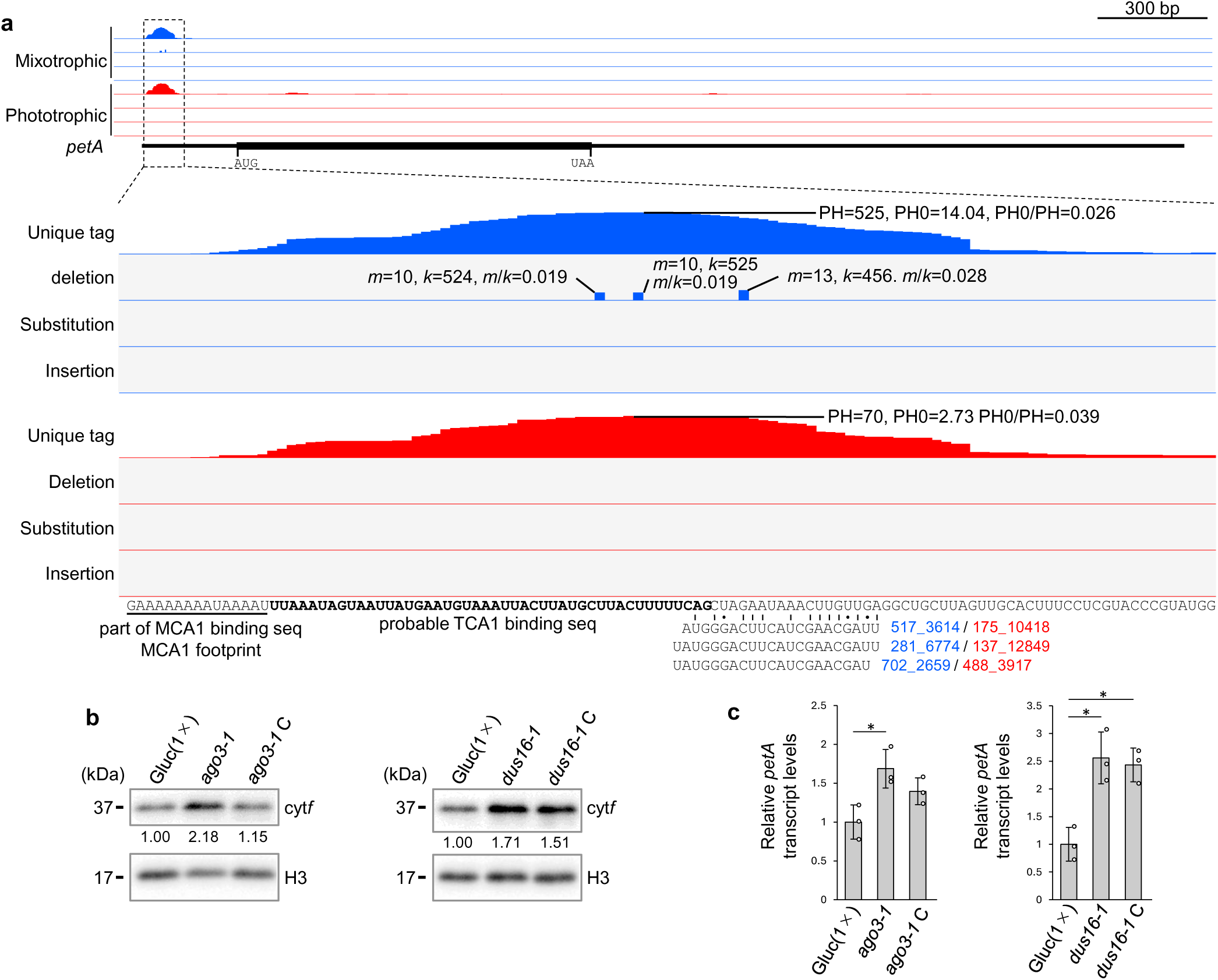
AGO3–RISC binds to the 5’ UTR of the *petA* transcript, and increased cyt*f* protein levels are present in the *ago3* and *dus16* mutants. (a) Location of unique tag clusters and CIMS on the *petA* transcript. The sequences below show regions with the footprint for the M-factor MCA1 (underlined) and for the T-factor TCA1 (in bold). (b) Immunoblot analysis of cyt*f* protein abundance in the *ago3-1* and *dus16-1* mutants and their partly complemented strains *ago3-1 C* and *dus16-1 C*. Protein samples were collected at ZT8. cyt*f* and H3 protein levels were quantified from the signal intensity, and cyt*f* abundance was normalized to that of H3, with the values from Gluc(1×) set to 1. (c) RT-qPCR analysis of *petA* mRNA levels in the indicated strains. RNA samples were collected from synchronized cells at ZT6. *petA* transcript levels were normalized to those of *NUOP4*. The value corresponding to Gluc(1×) for *petA* levels was set to 1, and the other values were adjusted accordingly. The values presented are the means ± SD of three independent biological replicates. A two-tailed, unpaired t-test was employed to identify statistically significant differences. Asterisks indicate statistically significant differences (P < 0.05).

The binding sequence for the translation-accelerating factor TCA1 (translation of cytochrome *b_6_f* complex *petA* mRNA)^32,33^ was located within the *petA* read clusters (Fig. 5), while the binding sequence for the mRNA-stabilizing factor MAC1 (maturation of psaC 1)^34^ was present in the *psaC* read clusters (Supplementary Fig. 9). Both TCA1 and MAC1 are positive regulators of protein production, and their binding sequences overlap with complementary sRNA sequences (Fig. 5, Supplementary Fig. 9). AGO3–RISC and M/F factors appear to compete for binding to chloroplast RNAs, which may lead to reduced RNA stability and/or translation efficiency.

### sRNAs containing the loop region of their precursor are bound to AGO3

Mapping of sRNA and target reads to the genome with no size selection applied revealed an ∼43-nt stem-loop RNA resulting from cropping of the hairpin loop derived from the Cre13.g585175 locus (Fig. 6). We also detected a ∼31-nt 3’ arm-cleaved stem-loop RNA, losing about 12 nt from its 3’ end, with the 3’ end corresponding to bases 10–11 of the 5’ arm, indicating cleavage by AGO3 (Fig. 6c). Additionally, we detected nucleotides 20–26 corresponding to the 5’ arm–loop RNA, likely produced similarly to Ago2-cleaved pre-miR-451 in vertebrates^35,36^. This observation suggests that the ∼43-nt stem-loop RNA is incorporated into AGO3, with the 3’ arm cleaved by slicer activity and subsequent exonuclease trimming from the cleavage site to the loop. By contrast, the 3’ arm sRNA variants were not aligned at both ends (Fig. 6c). This stem-loop RNA likely results from DCL3-induced cropping, but the stem length (17 nt with a single mismatch) is too short to remove the loop. Consequently, the incorporation of the stem-loop RNA into AGO3 produces a 3’ arm-cleaved stem-loop RNA through AGO3 cleavage and a 5’ arm-loop RNA via subsequent exonuclease activity. Alternatively, the 3’ arm sRNA may arise from the endonuclease-mediated cleavage of the loop in the stem-loop RNA.

**Fig. 6.**
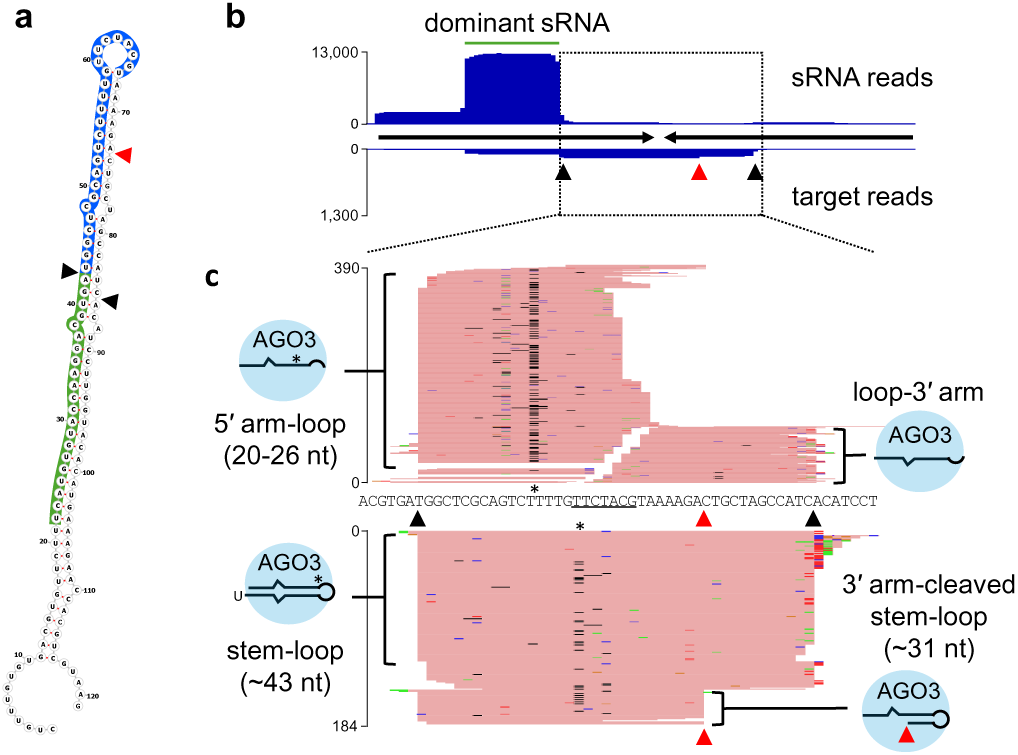
A non-canonical sRNA, similar to vertebrate pre-miR-451, is produced from Cre13.g585175. (a) Predicted structure of the sRNA precursor produced from Cre13.g585175. The sequence highlighted in green represents the dominant sRNA produced from the precursor. The sequences highlighted in blue correspond to sRNAs containing loop regions, termed 5’ arm-loop RNAs. Black triangles mark cleavage points likely processed by DCL3, while red triangles indicate cleavage points potentially processed by AGO3. (b) Mapping of sRNA and target reads to the inverted repeat sequence of the precursor shown in (a). Opposing black arrows in the center represent inverted repeats, with sRNA reads mapped above and target reads below. All deduplicated reads from each library, regardless of length, were used for mapping. The green line indicates the dominant sRNA produced from the precursor shown in (a). The black and red triangles are as in (a). The *y*-axis represents the number of deduplicated reads. (c) Enlarged view of the central part of the inverted repeat region outlined by the dashed box in (b). The sequence of the expanded region is shown in the center, with the loop region underlined. Pink horizontal lines indicate reads mapped to the sense strand, with the accumulation of sRNA library reads at the top and target library reads at the bottom. The black and red triangles are as in (a). The *y*-axis represents the number of deduplicated reads. Mutations in the reads are color-coded: black, deletion; red, mutation to T; green, mutation to A; blue, mutation to C; yellow, mutation to G. Asterisks (*) indicate probable sites of high-frequency crosslinking with AGO3.

We similarly identified a ∼40-nt stem-loop RNA in the precursor produced by Cre04.g229050, likely generated by DCL3 cropping (Supplementary Fig. 10). Unlike in the previous case, we did not detect a 3’ arm-cleaved stem-loop RNA; instead, we observed 5’ arm-loop intermediates and a 5’ arm-loop RNA, possibly produced by trimming of the 3’ end from the stem-loop RNA (Supplementary Fig. 10). The high frequency of untemplated U residues at the 3’ ends of the stem-loop and loop-3’ arm RNA suggests active degradation from this end^37^.

In *Chlamydomonas*, we detected both the 5’ arm-loop RNA and its precursor, the stem-loop RNA, using HITS-CLIP, but we only clearly identified two sRNA loci as sources of these RNAs. The presence of deletion clusters indicates that 5’ arm-loop RNAs are likely bound to AGO3, but their specific role in repressing gene expression remains unclear.

## Discussion

In this study, we identified 133 putative candidate AGO3 target genes. Notably, the transcript levels of two of these target genes, *CAS* and *petA*, were elevated in the *ago3-1* and *dus16-1* mutants, suggesting that AGO3–RISC likely represses the expression of these genes. Further investigation is required to understand how their repression affects the biology of *Chlamydomonas*. Knocking out the sRNAs targeting these transcripts may be necessary for deeper insights. We attempted to detect the protein abundance corresponding to other candidate genes but faced challenges due either to a lack of specific signals when a commercial antibody was available or to the unavailability of specific antibodies. To fully explore the role of sRNAs in *Chlamydomonas*, it will be crucial to determine whether the expression of the other target genes is similarly repressed and to uncover the physiological implications of this repression. The next step will be mapping the complete AGO3–mRNA interactome under varying growth conditions, for example over the course of the cell cycle, over the life cycle, and under stress conditions, to reveal how the suppression of gene expression relates to biological phenomena.

Three transcripts (Cre12.g557050.t1.2, Cre11.g467707.t1.1, and Cre01.g042750.t1.2), previously detected using degradome-seq^38^, were among the 1,213 genes with *k* ≥ 1 and *m*/*k* > 0 CIMS (Supplementary Table 16); however, the absence of CIMS and unique tag clusters at predicted miRNA-binding sites raises concerns about potential false positives. Among three additional transcripts (Cre17.g697550.t1.1, Cre17.g713200.t1.1, and Cre13.g608000.t1.2) confirmed to be cleaved by AGO via the 5’ rapid amplification of cDNA ends (5’ RACE)^14,21^, only Cre17.g697550.t1.1 had six unique tags at the cleavage site under phototrophic conditions, but we detected no CIMS. *PHOT* and Cre16.g683650, whose encoded proteins showed increased abundance in mutants defective in miRNA production^16,21^, were also not included in the 131 identified genes (Supplementary Table 15). Among the 131 genes, only *RIBOSOMAL PROTEIN L 22* (*RPL22*, Cre07.g357850) was previously confirmed as a target of miRNA-mediated translation inhibition in a complex omics analysis^22^. The 1,213 genes with CIMS that did not meet our stringent criteria included 10 previously identified targets, but were likely not significantly enriched. These discrepancies may stem from effects of the varying culture conditions on miRNA functionality, which is responsive to the cell cycle and stress. Moreover, the detection of target mRNAs by AGO3 HITS-CLIP depends on their absolute levels associated with AGO3–RISC. To comprehensively identify AGO3–RISC target genes in *Chlamydomonas*, improving HITS-CLIP sensitivity and analyzing diverse culture conditions will be essential.

Many chloroplast mRNAs are stabilized or their translation is facilitated by transcript-specific RNA-binding proteins, called maturation/stabilization factors (M factors) and translation factors (T factors). These proteins protect mRNAs from degradation, leading to the detection of small RNA footprints (15–41 nt) in the binding regions^33^. Notably, the unique tag cluster and CIMS positions on the *petA* and *psaC* transcripts overlapped with the binding sites of TCA1 (a T factor) and MAC1 (an M factor), respectively (Fig. 5, Supplementary Fig. 9). The increased *petA* mRNA and cyt*f* protein levels in the *ago3-1* mutant (Fig. 5) suggest that AGO3 represses *petA* expression, although insufficient complementarity would prevent cleavage by AGO3–RISC, if AGO3 is present in the chloroplast. This result suggests that AGO3–RISC may affect *petA* mRNA stability and/or translation efficiency without slicing.

While a chloroplast localization for AGO3 is unconfirmed, the overlap of its binding region with TCA1 suggests that AGO3 and TCA1 may compete for the *petA* mRNA, influencing its translation. In future research, including studies of AGO3 localization, mutation analyses of the *petA* and *psaC* binding sites and the knockout of the sRNA genes are needed to validate this model.

We have established a reference protocol for a HITS-CLIP analysis of *Chlamydomonas*. The PAR-CLIP method, developed after HITS-CLIP, enhances crosslinking efficiency by using photoreactive nucleosides and UV light irradiation, improving detection sensitivity^39^; however, nucleosides like 4SU and 6SG are toxic and reactive, complicating RBP–RNA detection in photosynthetic organisms. By contrast, HITS-CLIP can freeze RBP–RNA interactions under any conditions, making it suitable for studying RBPs in *Chlamydomonas*. Recent genome editing advancements should allow HITS-CLIP to target all nucleus-encoded RBPs, promising insight into RBP-related biological phenomena.

## Methods

### Algal strains

The *Chlamydomonas reinhardtii* strain CC-124 was obtained from the Chlamydomonas Center (https://www.chlamycollection.org/). The *ago3-1* mutant was isolated by random insertional mutation into a Gluc(1×) strain expressing a luciferase reporter construct carrying a sequence fully complementary to miR9897-3p^13^. The *ago3-1* C strain is a genetic complementation strain obtained by introducing the *AGO3* gene cloned in the expression vector p3xHA-2xgp62-3xFLAG(N) into *ago3-1*^13^. The *dus16-1* mutant was isolated in the same way as *ago3-1*^15^. The strain *dus16-1* C is a genetic complementation line of *dus16-1*^15^.

### Culture conditions

The *ago3-1* C strain used for the HITS-CLIP analysis was grown in Tris-acetate phosphate (TAP) medium^40^ for the mixotrophic culture or in high-salt medium (HSM)^41^ for phototrophic culture in an Erlenmeyer flask, with swirling at 120 rpm at 25°C under continuous white light at ∼150 µmol/m^2^/s. Culture synchronization was performed in HSM for at least two weeks under a 12-h light/12-h dark photoperiod (∼80 µmol/m^2^/s white LEDs) with a 25°C/18°C day/night cycle. The photosynthetic photon flux density values were measured with a light analyzer LA-105 (Nippon Medical & Chemical Instruments). The cell titer in the culture medium was measured using a Countess II FL automated cell counter (Thermo Fisher Scientific).

### Construction of AGO3 HITS-CLIP libraries

An overview of the procedure is shown in Supplementary Fig. 1. All steps were optimized with reference to Moore et al.^25^ and are explained below:

**UV irradiation and crosslinking:** Cultures in mid-log phase (2–3 × 10^6^ cells/ml) were chilled on ice for 5 min and then centrifuged at 2,000 *g* for 3 min at 4°C to collect the cells. The cell pellets were suspended in ice-cold medium (TAP or HSM) at a cell density of 2.5 × 10^7^ cells/ml. A 40-ml aliquot of the cell suspension (corresponding to 1 × 10^9^ cells) was poured into a 25 × 15 cm square glass dish on ice and placed in a UV crosslinker (FS-1500, Funakoshi). The suspension was irradiated with UV light (254 nm) at energies of 300, 300, 200, and 200 mJ/cm^2^, giving a total energy of 1,000 mJ/cm^2^. Between each exposure, the glass dish was gently shaken several times to mix the cells and ensure that each cell was exposed to UV light as evenly as possible.

**Immunopurification:** The UV-irradiated cell suspension was collected from the glass dish into centrifuge bottles and centrifuged at 2,000 *g* for 3 min at 4°C. The cell pellet was washed once with phosphate-buffered saline (PBS) before being resuspended in 10 ml PBS containing 25 µl protease inhibitor cocktail for plants (P9599, Merck). To lyse the cells, the concentrated cell suspension was ejected from an airbrush (PS-267 0.2 mm, GSI Creos) at a pressure of 0.7 MPa of nitrogen gas into a 50-ml tube^42^. Two ejections were carried out, and microscopy observations confirmed that almost 100% of the cells had been destroyed. The cell lysate was mixed with 25% (v/v) NP-40 and 25% (v/v) Triton X-100 to a final concentration of 0.5% (v/v) each, mixed well, and allowed to stand on ice for 10 min. The cell lysate was centrifuged at 16,000 *g* for 10 min at 4°C and the entire supernatant (called S16 below) was collected. A 50-µl aliquot (equivalent to 1.5 mg) Dynabeads Protein G (10004D, Thermo Fisher Scientific) was washed twice with PBS-NT (PBS containing 0.5% (v/v) NP-40, 0.5% (v/v) Triton X-100), resuspended in 150 µl PBS-NT, and rotated with 12.5 µl (12.5 µg) FLAG M2 antibody (F1804, Merck) for 30 min at room temperature. The beads were then washed twice with PBS-NT and resuspended in 100 µl PBS-NT to produce a FLAG bead solution. Unless otherwise stated below, DynaMag-2 (12321D, Thermo Fisher Scientific) was used for bead collection and washing. The FLAG beads solution was added to S16, rotated at 4°C for 30 min, and then washed three times in PBS-NT, once in 5× PBS-NT (5× PBS containing 0.5% (v/v) NP-40, 0.5% (v/v) Triton X-100), and once in PBS-NT.

**RNA fragmentation:** The beads remaining after the removal of PBS-NT were pre-warmed for 2 min at 22°C, and then suspended in 1 ml of RNase A solution (0.5 ng/ml RNase A (312-01931, Nippon Gene), 10 U/ml RQ1 DNase (M6101, Promega), 100 mM Tris-HCl pH 7.5, 10 mM MgCl_2_, 0.5% (v/v) NP-40, and 0.5% (v/v) Triton X-100), which was also pre-warmed to 22°C. The mixture was placed on a thermomixer (Eppendorf) and shaken at 1,000 rpm, with 15-s ON/60-s OFF cycles for 5 min at 22°C. The solution was then immediately cooled on ice and the RNase A solution was removed. The FLAG beads were washed three times with PBS-NT, twice with 5× PBS-NT, and twice with PNK buffer (100 mM Tris-HCl pH 7.5, 0.5% (v/v) NP-40, 0.5% (v/v) Triton X-100) to wash out the residual RNase A.

**Dephosphorylation:** The beads were resuspended in 30 µl of CIP solution (7.5 U of Quick CIP (M0525L, New England Biolabs) in 30 µl of 1× rCutSmart Buffer (New England Biolabs)) and incubated at 37°C for 20 min with shaking at 1,000 rpm, with 15-s ON/60-s OFF cycles, to dephosphorylate the 3’ end of the RNAs. The beads were washed once with PNK buffer, twice with PNK-EGTA (100 mM Tris-HCl pH 7.5, 100 mM EGTA, 0.5% (v/v) NP-40, 0.5% (v/v) Triton X-100), and twice with PNK buffer.

**3’ adaptor ligation and phosphorylation:** The beads were resuspended in 40 µl of ligation solution (2 µM CLIP-3 adaptor, 50 mM Tris-HCl pH 7.5, 10 mM MgCl_2_, 10 mM DTT, 2 mM ATP, 10 µg/µl bovine serum albumin (BSA), 25% (w/v) polyethylene glycol (PEG) 6000, and 0.25 U/µl T4 RNA ligase (EL0021, Thermo Fisher Scientific)) and incubated overnight at 15°C, with shaking at 1,000 rpm and 15-s ON/60-s OFF cycles. The CLIP-3 adaptor (P-CCCGAGCCCACGAGAC-DIG), an RNA oligo with its 5’ end phosphorylated and its 3’ end labeled with digoxigenin (DIG), was purified with high-performance liquid chromatography (HPLC) and used in the reaction. After the ligation reaction, the beads were washed once with PNK buffer and twice with 5× PBS-NT. They were then resuspended in 30 µl of PNK solution (70 mM Tris-HCl pH 7.6, 10 mM MgCl_2_, 5 mM DTT, 1 mM ATP, and 0.5 U/µl of T4 polynucleotide kinase (M0201S, New England Biolabs)) and incubated at 37°C for 20 min, with shaking at 1,000 rpm and 15-s ON/60-s OFF cycles, to phosphorylate the 5’ ends of RNAs. After the phosphorylation reaction, the beads were washed once with PNK buffer, once with 5× PBS-NT, and once with PNK buffer.

**PAGE separation:** The beads were resuspended in 20 µl of sample buffer (5 µl of 4× Bolt LDS sample buffer (B0007, Thermo Fisher Scientific), 14 µl of PNK buffer, and 1 µl of 1 M DTT) and incubated at 70°C for 10 min, with shaking at 1,000 rpm, to release AGO3 from the beads. The supernatant was divided into 1 µl (5%) and 19 µl (95%) and separated by electrophoresis using Bolt 8% (w/v) Bis-Tris Plus Gel (12 well, NW00082BOX, Thermo Fisher Scientific), 1× Bolt MOPS SDS Running Buffer (20×, B0001, Thermo Fisher Scientific), and a Mini Gel Tank (A25977, Thermo Fisher Scientific).

**Crosslinker detection (northwestern analysis):** After electrophoresis, the 5% sample lane was separated, and the resolved crosslinkers were transferred onto a polyvinylidene fluoride (PVDF) membrane (Immobilon-P, IPVH00010, Merck) using a Mini Blot Module (B1000, Thermo Fisher Scientific) and Bolt Transfer Buffer (BT0006, Thermo Fisher Scientific). The membrane was incubated with anti-DIG-AP-Fab fragments (11093274910, Merck), and the signals derived from DIG generated by the addition of SDP-star substrate (T2146, Thermo Fisher Scientific) were captured using a Multiimager II (BioTools).

**In-gel digestion and RNA purification:** After the presence of crosslinked complexes was confirmed by northwestern analysis, a gel slice was cut out from the 140–170-kDa range, where AGO3–RNA crosslinks were present, from the lane containing 95% of the sample. The gel slice was crushed into small fragments and frozen at −80°C for 30 min. The gel fragments were then thawed on ice and added to 100 µl of Proteinase K solution (100 mM Tris-HCl pH 7.5, 50 mM NaCl, 10 mM EDTA pH 8.0, and 4 µg/µl Proteinase K (169-21041, Fujifilm)) and incubated at 37°C for 120 min, with shaking at 1,000 rpm and 15-s ON/60-s OFF cycles to degrade the proteins, mainly FLAG-AGO3, in the gel. To this reaction solution, 100 µl of urea (7 M urea in 100 mM Tris-HCl pH 7.5, 50 mM NaCl, and 10 mM EDTA pH 8.0) was added, and the reaction was incubated for an additional 2 h under the same conditions to elute the RNA. The gel slurry was filtered through a Spin-X column (8162, Corning) to remove the gel fragments, and the eluate obtained was collected in DNA LoBind tubes (0030108051, Eppendorf), which were used for subsequent reactions. A volume of 23 µl of 5 M potassium acetate was added to the eluate, and the mixture was allowed to stand on ice for 30 min to precipitate the sodium dodecyl sulfate (SDS). The eluate was centrifuged at 5,000 *g* for 3 min at 4°C, and then 45 µl sodium acetate (pH 5.2), 3 µl GlycoBlue (AM9516, Thermo Fisher Scientific), and 400 µl isopropanol were added to the resulting supernatant for incubation at −30°C for 60 min to precipitate the RNAs. The solution was centrifuged at 16,000*g* for 30 min at 4°C, and the precipitated RNA was washed twice with 80% (v/v) ethanol.

**5’ adaptor ligation:** The RNA pellet was dissolved in 10 µl of 5’ ligation mix (2 µM 5’-CLIP-adaptor, 50 mM Tris-HCl pH 7.5, 10 mM MgCl_2_, 10 mM DTT, 2 mM ATP, 10 µg/µl BSA, 25% (w/v) PEG 6000, and 0.5 U/µl of T4 RNA ligase) and incubated overnight at 16°C to ligate the 5’ adaptor. An HPLC-purified RNA oligo, 5’-CLIP-adaptor (5’-CGUCGGCAGCGGUCAGAUGUGUGUAUAUAAGAGAGACAGNNNNNNNNN-3’) (Fasmac), was used. After the ligation reaction, 100 µl Milli-Q water, 20 µl sodium acetate (pH 5.2), 1 µl Glycoblue, and 200 µl isopropanol were added to the reaction solution and the mixture was cooled at −30°C for 60 min. The ligation products were precipitated by centrifugation at 16,000 *g* for 30 min at 4°C, washed twice with 80% (v/v) ethanol, air-dried, and dissolved in 10 µl of 98% formamide.

**Size selection (ligation products):** RNA dissolved in formamide was separated on 8% (w/v) acrylamide/7 M urea/1× TBE gels together with Low Range ssRNA Ladder (N0364S, New England Biolabs), stained with Diamond nucleic acid dye (H1181, Promega), and visualized using excitation light at ∼500 nm. A range of ligation products with fragmented mRNA (95–140 nt) and a range of ligation products with sRNA (75–90 nt) were cut out separately, crushed into small fragments, and frozen at −80°C for 30 min. Nucleic acid extraction buffer (300 mM sodium acetate pH 5.2, 100 mM Tris-HCl pH 7.5, and 1 mM EDTA pH 8.0) was added to the gel fragments and incubated at 22°C for 4 h, with shaking at 1,000 rpm and 15-s ON/60-s OFF cycles, to elute the RNA. The eluted RNA was filtered through a Spin-X column before being added to 1.5 µl Glycoblue and an equal volume of isopropanol, stored overnight at −30°C, and centrifuged at 16,000 *g* for 30 min at 4°C. The precipitated ligation product was washed twice with 80% (v/v) ethanol and dissolved in Milli-Q water.

**Reverse transcription:** The purified RNA was reverse transcribed using the CLIP-3 PCR primer (5’-GTCTCGTGGGCTCGG-3’) and SuperScript IV reverse transcriptase (18090010; Thermo Fisher Scientific). The first-strand cDNA was then subjected to PCR using the CLIP-5 PCR primer (5’-TCGTCGGGCAGCGTC-3’), CLIP-3 PCR primer, and KOD Plus Neo DNA polymerase (KOD-401, Toyobo). The reaction was stopped after 16 cycles to amplify the sRNA library and 18 cycles to amplify the fragmented target mRNA library.

**Library preparation:** The RT-PCR products were separated on 8% (w/v) acrylamide/1× TBE gels and visualized with Diamond nucleic acid dye. Gels in the range of 80–90 bp were excised from lanes in which the RT-PCR products of sRNA were separated. Gels in the range of 95–130 bp were excised from gels in which the RT-PCR products of fragmented target mRNAs were separated. The excised gels were crushed into small fragments and frozen at −80°C for 30 min. The gel fragments were added to 200 µl of nucleic acid extraction buffer and incubated at 22°C for 4 h, with shaking at 1,000 rpm with 15-s ON/60-s OFF cycles, to elute the DNA contained in the gel. To the eluted DNA filtered through a Spin-X column, 2 µl Glycoblue and 300 µl isopropanol were added, and the mixture was stored at −30°C overnight. The solution was centrifuged at 16,000 *g* for 30 min at 4°C, and the precipitated DNA was washed twice with 80% (v/v) ethanol.

**Sequencing:** The RT-PCR products recovered from the gel were dissolved in Milli-Q water and amplified by index PCR using IDT for Illumina DNA/RNA UD index primers (20027213, Illumina) and KOD Plus Neo DNA polymerase. The index PCR was stopped after 8–10 cycles and the library DNA was purified using AMPure XP (A63880, Beckman Coulter). The library was sequenced on an Illumina NovaSeq 6000 instrument (150-bp paired-ends) (sequenced by Relixa, Japan).

### Bioinformatics analysis of the HITS-CLIP libraries

The bioinformatics analysis of the reads obtained from the HITS-CLIP libraries was performed according to the CLIP Tool Kit (CTK) pipeline^26^ (https://zhanglab.c2b2.columbia.edu/index.php/Standard/BrdU-CLIP_data_analysis_using_CTK), using the reference *Chlamydomonas* genome, primary transcript sequences, and annotation files (the original file is available at Phytozome (https://phytozome-next.jgi.doe.gov/info/Creinhardtii_v5_6). An overview of the pipeline is shown in Supplementary Fig. 3. Raw reads were filtered using fastq_filter.pl to remove the reads with a score below Q30 in the first 33 bases for the sRNA reads and in the first 73 bases for the target reads. From the filtered reads, the adaptor sequences (5’-CCGAGCCCACGAGAC-3’) were trimmed using cutadapt^43^, resulting in reads consisting of a Unique Molecular Identifier (UMI) sequence (8N) and an insert sequence. The UMI-insert reads were deduplicated using tastq2collapse.pl and then UMI sequences were stripped using stripBarcode.pl to obtain deduplicated reads. The counting of the sRNA reads (Supplementary Tables 11 and 12), analysis of sRNA length distribution, and analysis of the 5’ end preference of sRNA (Fig. 1) were performed using seqkit (version 0.16.0)^44^ and fastx_collapser (0.0.14). Datasets of 20–22-nt sRNA redundant reads and 30–100-nt target redundant reads were generated using seqkit, mapped to the whole genome or the whole primary (major) transcript sequence, which includes the primary (major) mature transcript sequence of the nuclear genes and the chloroplast and mitochondrial genome sequences^45^, using bwa (version 0.7.17) (-n 0.06)^46^ and converted to SAM files^47^. In the next step, the default parameters were used to obtain the output from the BED and mutation files with parseAlignment.pl; to obtain unique tags with tag2collapse.pl; to obtain mutations in unique tags with joinWrapper.py; and to merge triplicate data and mutation files with bed2rgb.pl. Peak detection was performed using tag2peak.pl with the genome sequence as the reference, and half-peak height (halfPH) was calculated with default parameters. For the peak detection and halfPH calculation with reference to the transcripts, the value for valley seeking was adjusted from -p 0.05 to -p 0.001 to achieve an FDR < 0.001. The attribution of the detected peaks was determined with bed2annotation.pl. Deletions, substitutions, and insertions were detected using getMutationType.pl from the unique tag BED file integrated with the mutation file. Significant CIMS with FDR < 0.001 were then detected using CIMS.pl. The attribution of these CIMS was determined with bed2annotation.pl. BED files for unique tags and CIMS with FDR < 0.001 were converted to bedgraph for visualization in IGV in tag2profile.pl. The results of the analyses were visualized in IGV (version 2.17.4)^48^.

The secondary structure of the RNAs was predicted using RNAfold^49^ (http://rna.tbi.univie.ac.at/cgi-bin/RNAWebSuite/RNAfold.cgi).

### psRNAtarget analysis

The psRNAtarget analysis^50^ was performed as follows. The sequence used as the target query was expanded from each halfPH by 30 nt on either side of the halfPH in the BED file and extracted using bedtools. To obtain the sequences used as sRNA queries, UMI-sRNA reads were first obtained by trimming downstream from the adaptor sequences from filtered reads obtained from sRNA libraries. UMI-sRNA reads from biological replicates were integrated, and a deduplication, including UMI sequences, was performed using fastX_collapser. Only sRNA sequences obtained from the deduplicated sRNA sequences that were a perfect match to the *Chlamydomonas* genome (version 6.1, https://phytozome-next.jgi.doe.gov/info/CreinhardtiiCC_4532_v6_1) were extracted. Only sRNA sequences with more than 100 reads per million were selected for use as sRNA queries. The parameters of the psRNAtarget analysis were adjusted as follows: “seed” from 2 to 8, “penalty for other mismatches” from 1 to 0.5, “extra weight in seed region” from 1.5 to 4 and “# of mismatches allowed in seed region” from 2 to 1.

### Immunoblot analysis

*Chlamydomonas* cells for analysis were pelleted by centrifugation at 5,000 *g* for 5 min at 4°C and resuspended in 10 µl sample buffer (100 mM Tris-HCl pH 6.8, 50 mM DTT, 50% (v/v) glycerol, 4% (w/v) SDS, 0.02% (w/v) bromo phenol blue) per 1 × 10^6^ cells. Boiled protein samples were separated by SDS-PAGE and electroblotted onto an Immobilon-P (Merck) membrane. To detect CAS, an anti-CAS antibody^31^ was used at a 1:20,000 dilution. To detect cyt*f*, an anti-cyt*f* antibody (AS06 119; Agrisera) was used at a 1:20,000 dilution. To detect ATPB, an anti-ATPB antibody (AS05 085; Agrisera) was used at a 1:20,000 dilution. To detect histone H3, an anti-histone H3 antibody (ab1791; Abcam) was used at a 1:10,000 dilution. To detect FLAG-AGO3, FLAG M2 antibody was used at a 1:10,000 dilution. HRP-conjugated anti-rabbit IgG antibody (#7074, Cell Signaling Technology) and HRP-conjugated anti-mouse IgG antibody (#7076, Cell Signaling Technology) were used as secondary antibodies at a 1:20,000 dilution. The signal was detected using Multiimager II with ECL substrate EzWestLumi plus (2332637, Atto). The signal intensities were measured with ImageJ (https://imagej.net/ij/download.html) and utilized for a comparative quantitative analysis. All immunoblot assays were performed in biological triplicate, and the presented results are representative.

### RNA analysis

The total RNA was purified using TRI reagent (TR118, Molecular Research Center) and concentrations were measured with a NanoDrop 1000 (Thermo Fisher Scientific). The poly(A) length assay was performed using a Poly(A) Tail-Length Assay Kit (764551KT; Thermo Fisher Scientific). G/I tailing and reverse transcription were performed using 300 ng of purified RNA from HSM synchronized cultured cells collected at zeitgeber 8 (ZT8, with lights on being ZT0) as a template. Poly(A) tail PCR for *DII4* was performed with ACT1-polyA-F3 (5’-GGATGCAGGGCTCTTCATTC-3’) and a universal PCR reverse primer, and for *CAS* with CAS1-polyA-F1(5’-GAGTCGGAGAAACTCGGGTAGTG-3’) and a universal PCR reverse primer. For gene-specific PCR, ACT1-polyA-F3 and ACT1-polyA-R1 (5’-CTCATGTAAAAATGCTACACACGAAATGGTC-3’) were used for *DII4* and CAS1-polyA-F1 and CAS1-polyA-R1 (5’-CCGCTGACCGGGACAATTAGAG-3) for *CAS*. Poly(A) tail PCR amplicons were purified with a NucleoSpin Gel and PCR Clean-up kit (740609, Macherey-Nagel), separated on 1% (w/v) agarose/Tris-acetate EDTA (TAE) gels for ethidium bromide staining, and subjected to a Bioanalyzer (Agilent Technologies) analysis with a High Sensitivity DNA kit (5067-4626, Agilent Technologies).

For the first-strand cDNA synthesis for RT-qPCR, 300 ng of total RNA was used as a template with the ReverTra Ace cDNA Synthesis Kit (Toyobo). A quantitative PCR was performed using KOD SYBR qPCR Mix (Toyobo) on a StepOne Real-Time PCR System (Thermo Fisher Scientific). The primers used for RT-qPCR were as follows: CAS-F1 (5’-CCGTGGCGCTGTATTACCTCAG-3’) and CAS-R5 (5’-TCAATGTCCTGGGGGTTCTTCAG-3’) for *CAS*; petA-F1 (5’-GCTTAGCAGTAAGTCCAGCTCAAGC-3’) and petA-R1 (5’-GGTGGTGCTAATTCAAAACCTTCTGG-3’) for *petA*; NUOP4-F1 (5’-TCATGATTAACGACGATTACAAGTACACG-3’) and NUOP4-R1 (5’-CCTGCTGTTAAACTCTACTGCAGCAG-3’) for *NUOP4*.

## Accession numbers

The HITS-CLIP raw data have been deposited with links to BioProject accession number PRJDB12253 in the DDBJ BioProject database.

## Acknowledgments

We thank Prof. Jun Minagawa for the kind gift of the anti-CAS antibody. This work was supported by JSPS KAKENHI (grant numbers 16H06279 (PAGS), 16K18480, 17H05734, 19H04728, 19K06710, and 22K06268), the Nagase Science and Technology Foundation, the Institute for Fermentation (IFO), the Takeda Science Foundation, and a Grant-in-Aid for a Core Research Project from Kochi University.

## Author contributions

T.Y. conceived and led the project. S.M. and K.S. carried out the immunoblot analysis. T.Y., H.T., and K.S. conducted the bioinformatics analysis using reads from the HITS-CLIP libraries. T.Y. optimized the conditions for the HITS-CLIP library construction and performed the library construction, psRNAtarget analysis, and RNA analysis. T.Y. analyzed the data and wrote the manuscript.

## Data availability

All data supporting the findings of this study are available from the corresponding author upon request. Processed data are available in the supplementary materials.

## Completing interests

Authors declare no competing interests.

## Supplementary Materials

Supplementary Figs. 1–10

Supplementary Tables 1–16

Supplementary Data 1

