## Supplementary Figure 1-10 for "Global identification of AGO3–RNA interactions in *Chlamydomonas* reveals the small RNA–mediated regulation of nuclear and chloroplast gene expression"

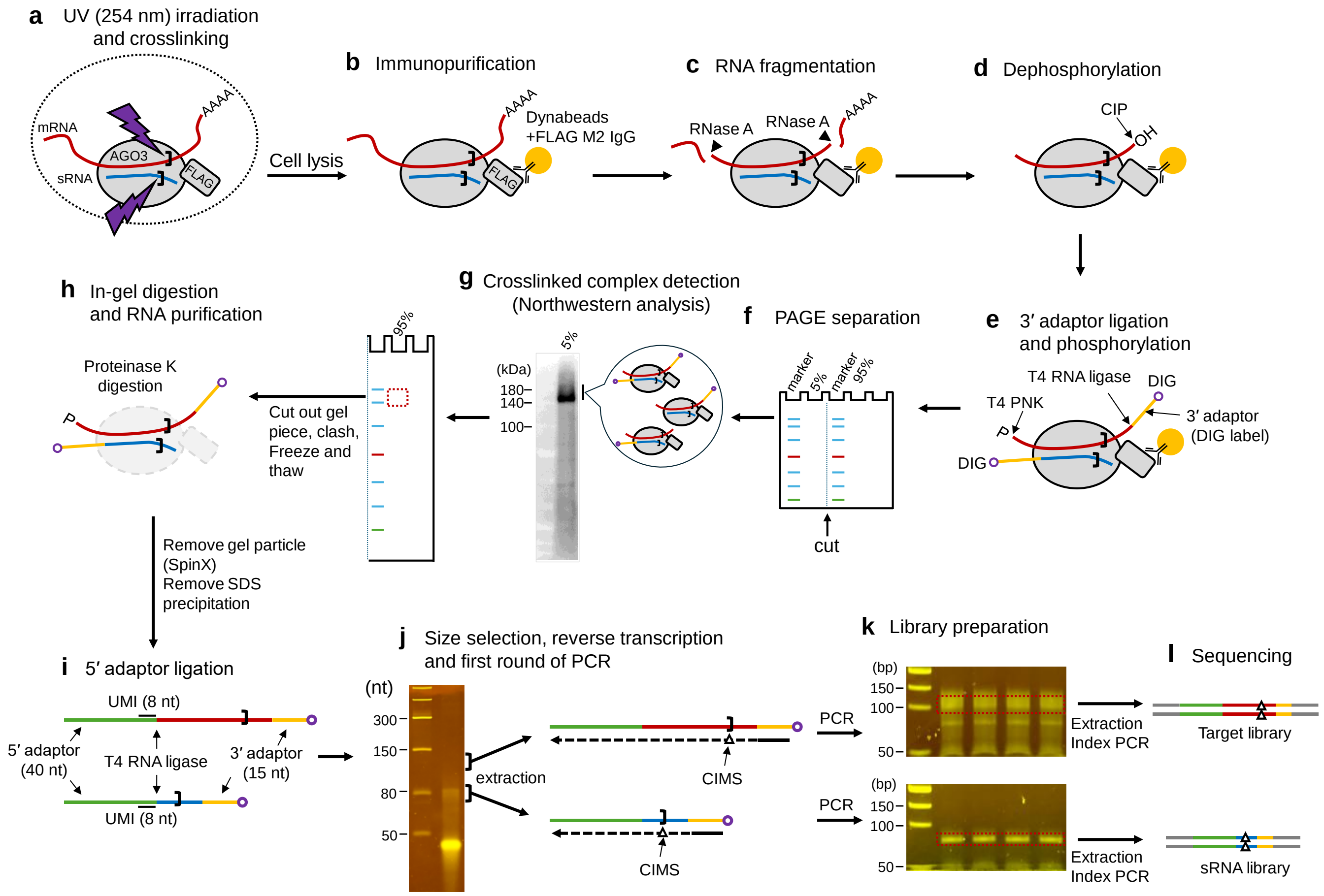


**Supplementary Figure 1. Overview of library preparation for the AGO3 HITS-CLIP analysis.**

(a) UV irradiation and crosslinking: *Chlamydomonas* cells are exposed to UV light, which induces crosslinking between proteins and RNAs.

(b) Immunopurification: The AGO3–RNA complexes are immunopurified using an anti-FLAG monoclonal antibody and Dynabeads Protein G incubated with the supernatants obtained by the centrifugation of the cell lysates after UV crosslinking.

(c) RNA fragmentation: The mRNAs crosslinked to AGO3 are fragmented by incubation with very low concentrations of RNase A.

(d) Dephosphorylation: The 3' end of the fragmented mRNA is dephosphorylated by incubation with calf intestinal alkaline phosphatase.

(e) 3' adaptor ligation and phosphorylation: 3' adaptors are ligated to the fragmented mRNAs and sRNAs, and the 5' ends of the RNA fragments and sRNAs are phosphorylated. The 3’ end of the 3' adaptor is labeled with digoxigenin (DIG) to enable detection and prevent self-concatemerization. All reactions up to this point are performed on beads.

(f) PAGE separation: The eluates from the Dynabeads are separated by PAGE, split into two fractions corresponding to 5% and 95% of the eluate. The 5% fraction is transferred to a PVDF membrane for the northwestern analysis, while the remaining 95% fraction is used for RNA extraction from the gel.

(g) Crosslinked complex detection: The AGO3–RNA crosslinked complexes ligated with a 3' adaptor are detected at a slightly higher molecular weight position than FLAG-AGO3 alone (~130 kDa).

(h) In-gel digestion and RNA purification: The FLAG-AGO3 protein is digested with proteinase K in-gel, and the crosslinked RNAs are extracted and purified.

(i) 5’ adaptor ligation: A unique molecular identifier (UMI) sequence (eight consecutive Ns) is added to the 3' end of the 5' adaptor.

(j) Size selection, reverse transcription, and first round of PCR: Ligation products with fragmented target mRNA and sRNA inserts are separated by PAGE. RNA is extracted from the gels, followed by reverse transcription and a first round of PCR.

(k) Library preparation: The PCR products are separated by PAGE, and DNA corresponding to the size of the sRNA and target libraries is extracted and used as a template for the index PCR.

(l) Sequencing: Short reads are obtained from the target and sRNA libraries.

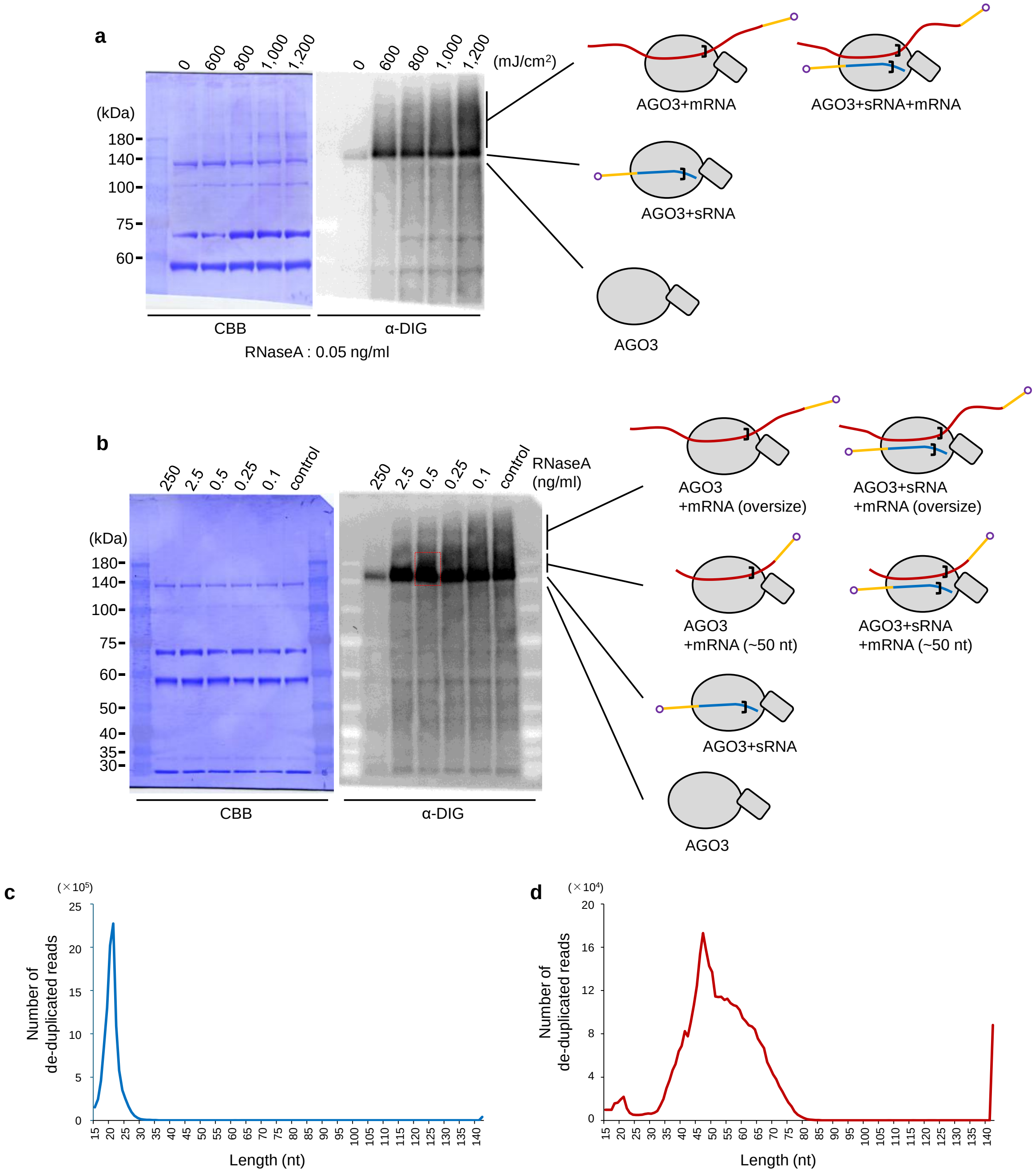


**Supplementary Figure 2. Total UV energy and concentrations of RNase A were optimized for the AGO3 HITS-CLIP method.**

(a) Northwestern analysis of immunopurified FLAG-AGO3 (Supplementary Fig. 1g) on cells irradiated with different total energy doses of UV light. Fragmentation was performed using 0.05 ng/ml RNase A. After detecting the DIG signal, the membrane was stained with Coomassie Brilliant Blue (CBB).

(b) Northwestern analysis of immunopurified FLAG-AGO3 treated with different concentrations of RNase A. AGO3–RNA crosslinked complexes from cells irradiated with 1,000 mJ/cm$^2$ UV were treated with the indicated concentrations of RNase A. After detecting the DIG signal, the membrane was stained with CBB.

(c, d) Distribution of the insert lengths (deduplicated) for the sRNA (c) and target (d) libraries.

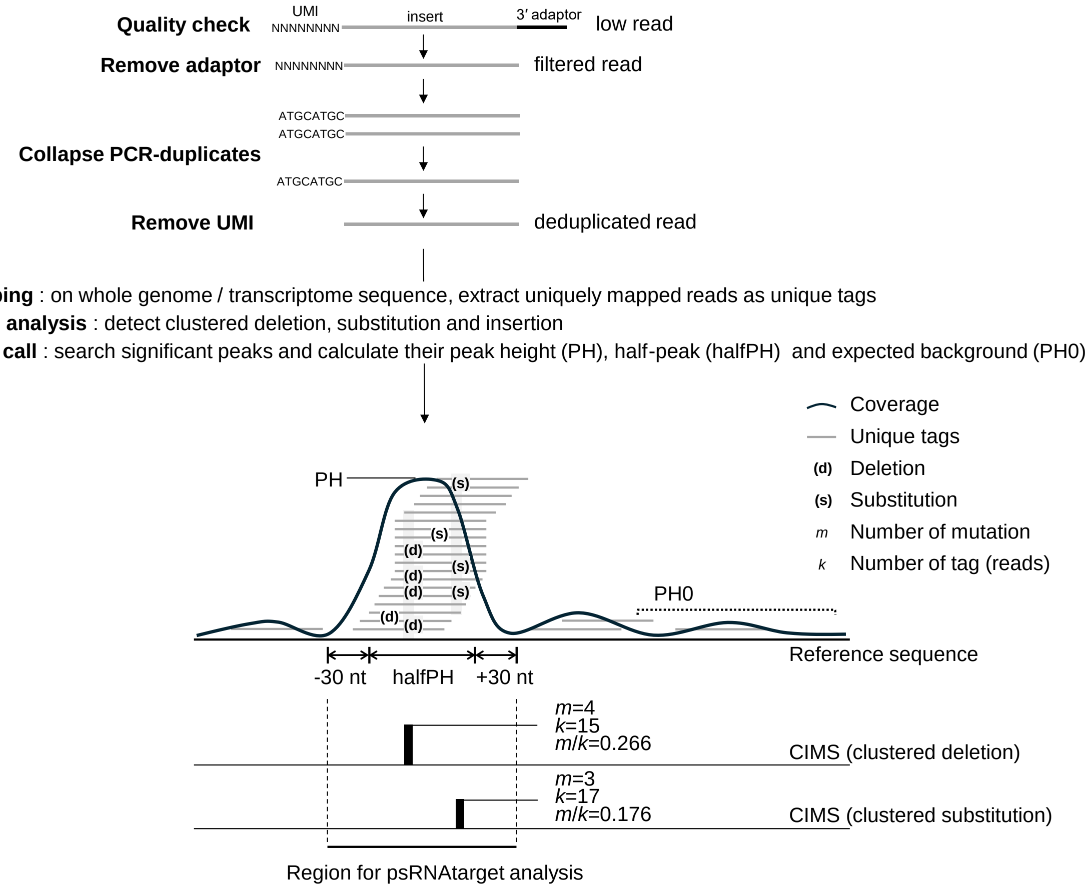


**Supplementary Figure 3. Overview of the bioinformatics analysis of AGO3-crosslinked RNAs**

Raw reads are filtered, their 3' adaptor sequences removed, and the resulting clean reads are deduplicated using UMI sequences to obtain deduplicated reads. These reads are mapped to reference sequences to obtain unique tags, which are then analyzed using the CTK toolkit pipeline to detect CIMS and peak calls. High-confidence CIMS are selected from the detected CIMS according to their *k*-value ($k \geq 100$) and *m*/*k*-value ($0.95 > m/k \geq 0.08$) (*m*, number of mutations; *k*, number of unique tags covering the mutation site). High-confidence peaks are selected from the detected peaks according to their PH value (PH ≥ 100) and PH0/PH value (PH0/PH ≤ 0.1). The sequence encompassing halfPH and 30 nucleotides on either side of the high-confidence peaks are considered as the region covered by each unique tag cluster, and peaks with high-confidence CIMS in this region are selected. Concurrently, peaks with sequences complementary to sRNAs within the halfPH ± 30 nt sequence are selected using psRNAtarget analysis. Transcripts with high-confidence peaks (high-confidence unique tag clusters) selected up to this point are considered candidate transcripts to which the AGO3–RISC binds.

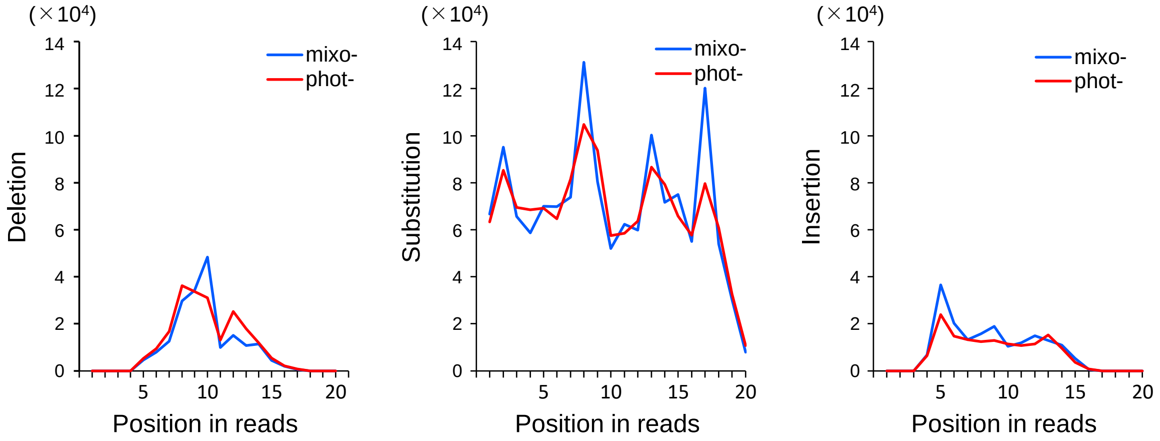


**Supplementary Figure 4. Positional aspects of deletions, substitutions, and insertions from the 5' end in deduplicated small RNA reads resemble those of mouse AGO2 in a HITS-CLIP analysis.**

The *x*-axis of each graph indicates the position from the 5' end of the sRNA. The *y*-axis shows the number of each type of mutation detected in all redundant reads, not limited to mutations in positions detected as CIMS.

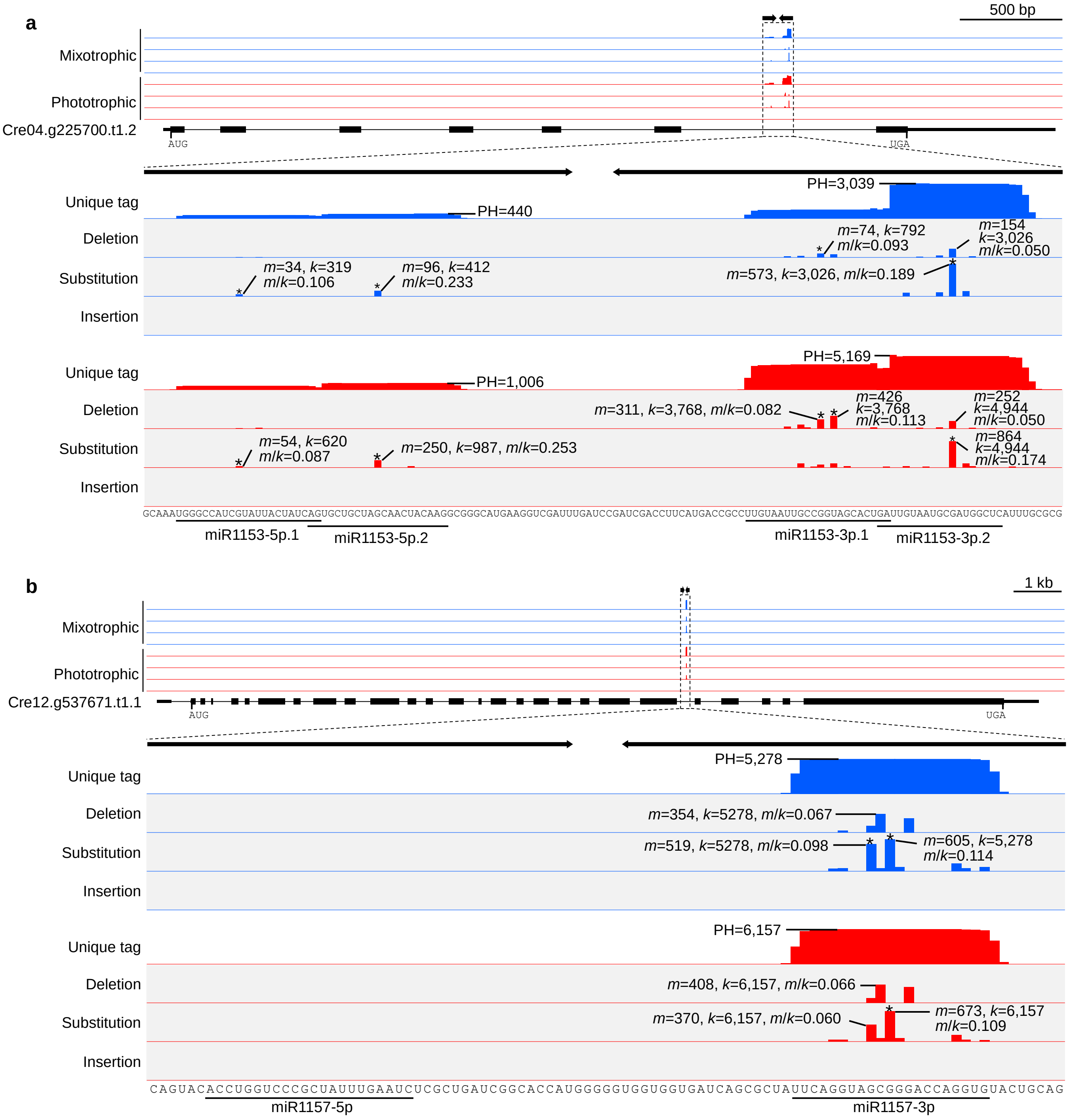


**Supplementary Figure 5. Cluster of unique sRNA tags and CIMSs with high *m*/*k* rates found in the *MIR1153* and *MIR1157* loci.**

An overview of Cre04.g225700.t1.2 (a), where the primary transcript of *MIR1153* is located, and of Cre12.g537671.t1.1 (b), where the primary transcript of *MIR1157* is located. The inverted repeat regions for each primary *MIR* transcript are highlighted with a dashed box. An enlarged view of this region is presented in the lower panels. The sequence of this region is shown at the bottom, with the sequence of the mature miRNAs underlined. PH, peak height; $m$, number of mutations; $k$, number of unique tags covering the mutation site. Each asterisk (*) marks CIMS with $m/k \geq 0.08$.

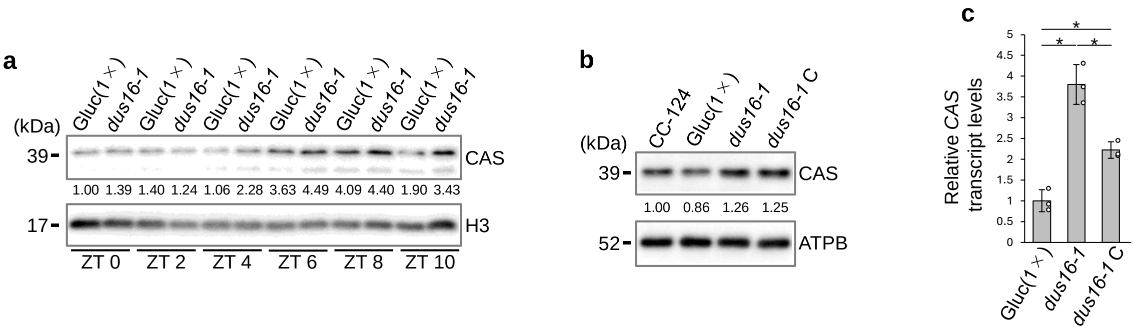


**Supplementary Figure 6. CAS protein abundance and *CAS* mRNA levels increased in the *dus16* mutant.**

(a) Immunoblot analysis of CAS protein abundance in Gluc(1×) and *dus16-1*. *dus16-1* is a mutant of *DUS16* isolated from the parental strain Gluc(1×). The two strains were grown synchronously under a 12-h light/12-h dark photoperiod and protein samples were collected every 2 h from the start of the light period (zeigeber time 0 (ZT0)) until 10 h after lights on (ZT10). Histone H3 was used as the loading control.

(b) Immunoblot analysis of CAS protein abundance in the *dus16-1* genetic complementation strain. The *dus16-1* C strain is a genetic complementation line expressing *DUS16-FLAG* in the *dus16-1* background. ATPB was used as loading control.

(c) RT-qPCR analysis of *CAS* mRNA levels in the indicated strains. RNA samples were collected from synchronized cells at ZT6. Asterisks indicate statistically significant differences ($P < 0.05$).

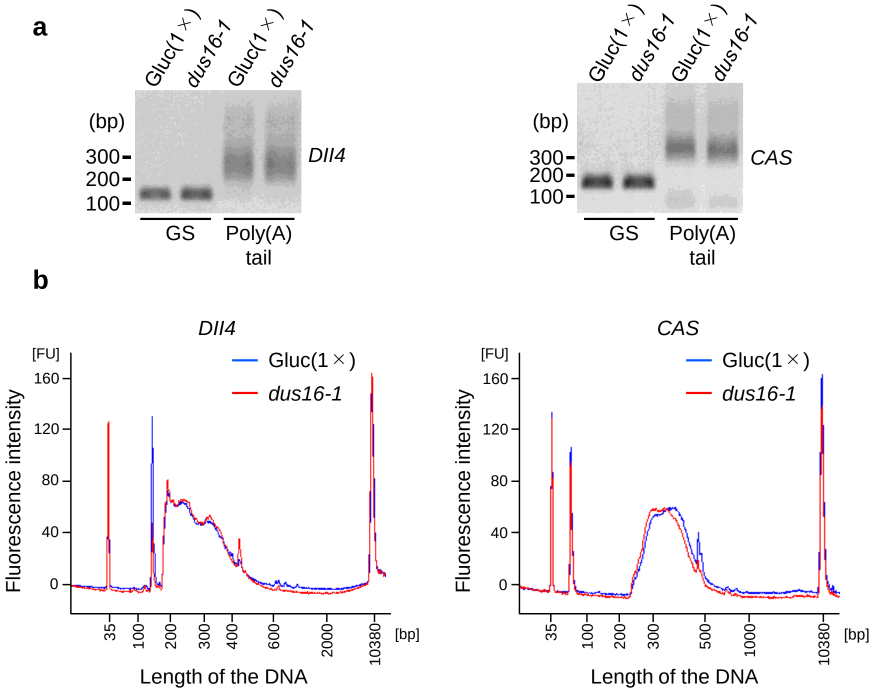


**Supplementary Figure 7. Poly(A) tail length of *CAS* mRNA is slightly longer in *dus16-1* than in its parental strain.**

(a) Agarose gel electrophoresis analysis of gene-specific PCR and poly(A) tail PCR products for the actin gene *Dll4* (also reported as *ACT1*, Cre13.g603700) and *CAS*. gs, gene-specific PCR amplicon; poly(A) tail, poly(A) tail PCR amplicon. All Poly(A) tail length assays were performed in biological triplicates and the presented results are representative.

(b) Bioanalyzer analysis of *Dll4* and *CAS* poly(A) tail PCR products.

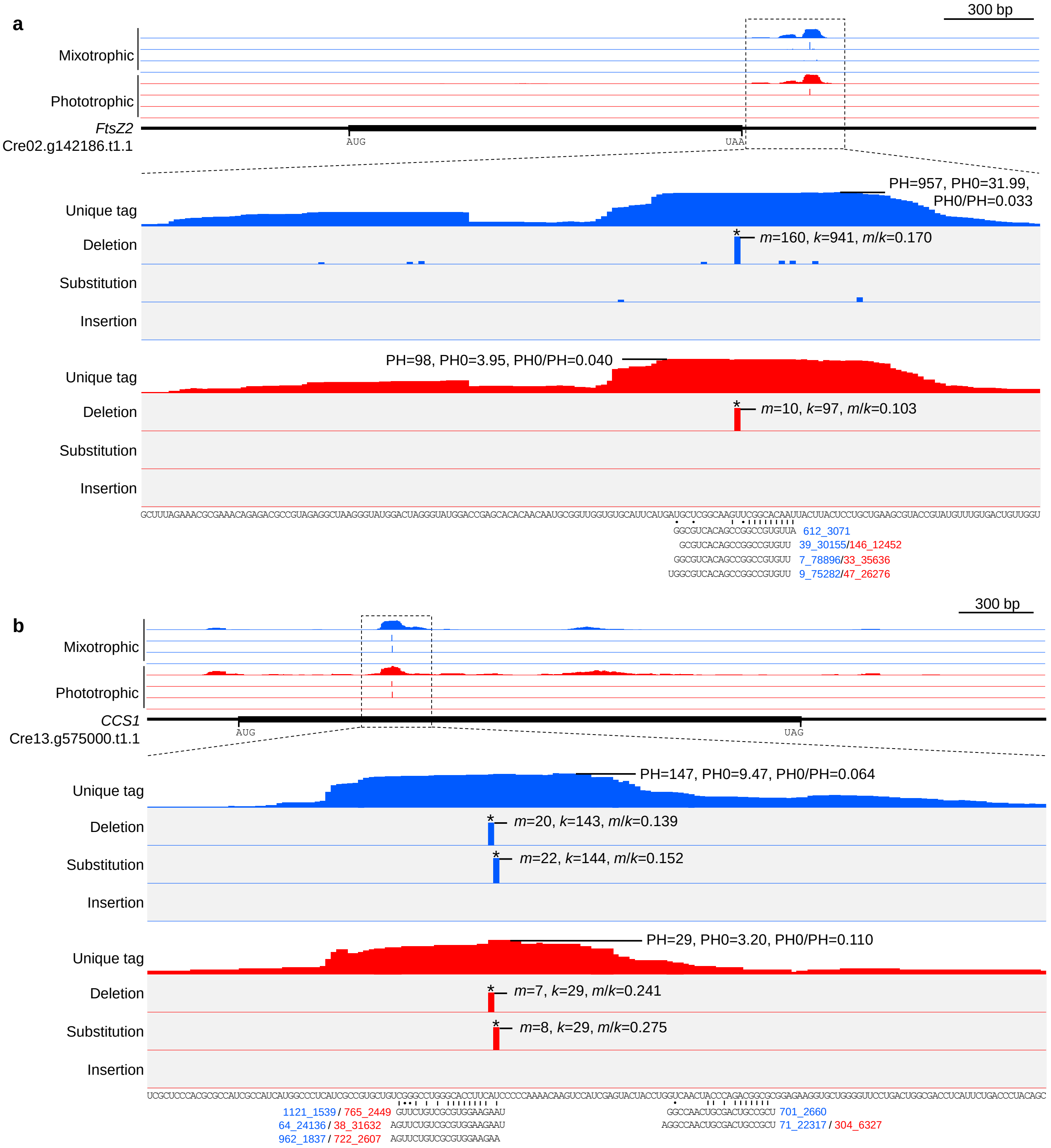


**Supplementary Figure 8. Unique tag clusters and CIMSs found in the *FtsZ2* and *CCS1* mRNAs.**

(a) represents *FtsZ2* (Creg.142186.t1.1) and (b) represents *CCS1* (Cre13.g575000.t1.1).

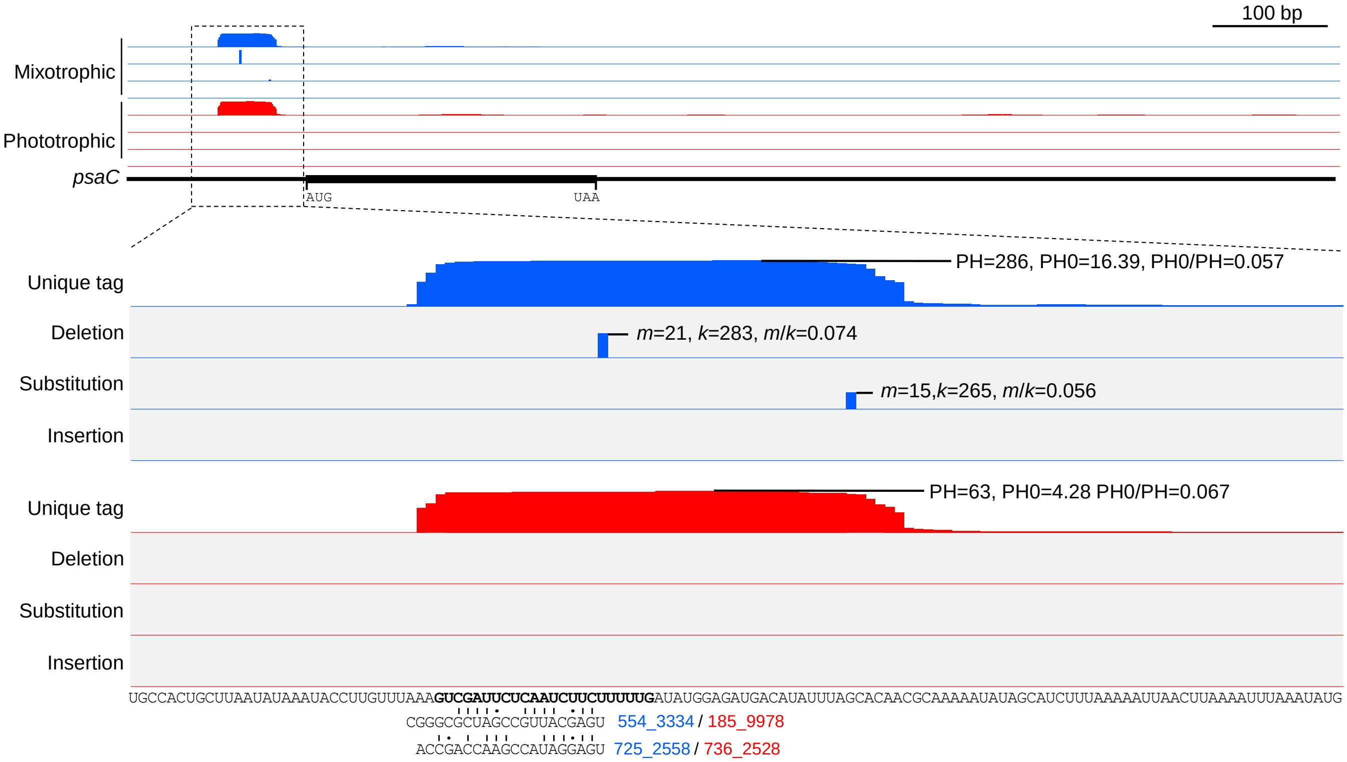


**Supplementary Figure 9. Unique tag clusters and CIMSs found in the *psaC* transcript.** In the sequences at the bottom, the sequence corresponding to the footprint of the T-factor MAC1 are in bold.

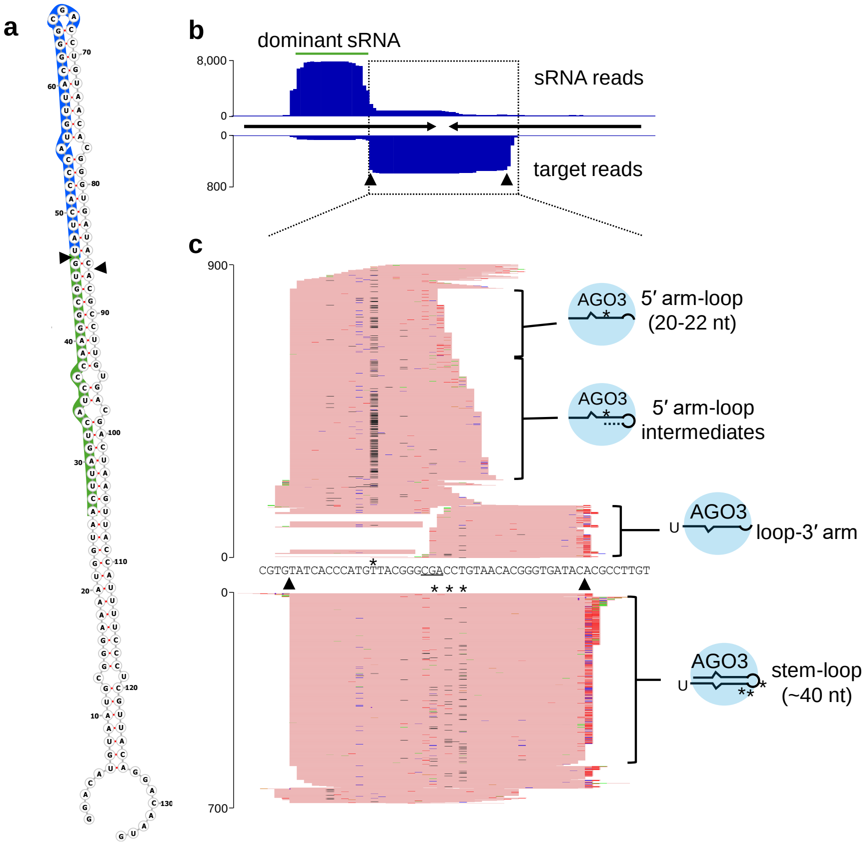


**Supplementary Figure 10. The sRNA precursor structure produced by Cre04.g229050, and the mapping of sRNA and target reads onto the precursor region.**

(a) Structure of the sRNA precursor produced from the Cre04.g229050 locus.

(b) sRNA and target reads mapping to the inverted repeat sequence of the precursor indicated in (a).

(c) Magnified view of the central part of the inverted repeat region shown in the dashed box in (b).
